# The second messenger c-di-AMP controls natural competence via ComFB signaling protein

**DOI:** 10.1101/2023.11.27.568819

**Authors:** Sherihan Samir, Sofía Doello, Andreas M. Enkerlin, Erik Zimmer, Michael Haffner, Teresa Müller, Lisa Dengler, Stilianos P. Lambidis, Shamphavi Sivabalasarma, Sonja-Verena Albers, Khaled A. Selim

**Affiliations:** Interfaculty Institute of Microbiology and Infection Medicine, Organismic Interactions Department, Cluster of Excellence “Controlling Microbes to Fight Infections”, Tübingen University, 72076 Tübingen, Germany; Microbial Biochemistry Group, Institute of Phototrophic Microbiology, Heinrich-Heine University Düsseldorf, 40225 Düsseldorf, Germany; Molecular Biology of Archaea, Institute of Biology, University of Freiburg, Freiburg, Germany; Microbiology Department, Faculty of Science, Ain Shams University, Cairo, Egypt; Signalling Research Centres BIOSS and/or CIBBS, Faculty of Biology, Freiburg University, Freiburg, Germany

## Abstract

Natural competence requires a contractile pilus system. Here, we provide evidence that the pilus biogenesis and natural competence in cyanobacteria are regulated by the second messenger c-di-AMP. Furthermore, we show that the ComFB signaling protein is a novel c-di-AMP-receptor protein, widespread in bacterial phyla, and required for pilus biogenesis and DNA uptake.

---

Second messengers are small molecules involved in regulating many processes in all kinds of organisms (Yoon & Waters 2021). Cyclic di-AMP is one of the recently discovered di-nucleotide-type second messengers only present in prokaryotes (Stülke & Krüger 2020; He et al. 2020; Yin et al. 2020; Mantovani et al. 2023a). Its functions have been mainly studied in firmicutes, where it plays an important role in osmo-adaptation by controlling potassium homeostasis and influencing transcription of genes related to osmoregulation and cell wall metabolism (Stülke & Krüger 2020; He et al. 2020; Yin et al. 2020; Herzberg et al. 2023; Nelson et al. 2013; Ren & Patel 2014; Foster et al. 2024). In cyanobacteria, c-di-AMP seems to control additional processes, including the diurnal metabolism via its binding to the carbon control protein SbtB to regulate glycogen metabolism (Rubin et al. 2018; Selim et al. 2021a). Although important roles for c-di-AMP have been acknowledged since its discovery, recent studies suggest broader regulatory impacts of c-di-AMP signaling with further functions yet to be elucidated (Mantovani et al. 2022; Haffner et al. 2023b). For instance, a role for c-di-AMP in controlling natural competence has been speculated in *Streptococcus pneumoniae*, although the molecular mechanism remained elusive (Zarrella et al. 2020). In this study, we aimed to investigate the involvement of c-di-AMP in natural competence.

Natural competence is a conserved mechanism of horizontal gene transfer that permits massive genetic variation and genomic plasticity via uptake of extracellular DNA, and it is the main cause of spreading antibiotic resistance and acquisition of virulence factors (Gibson & Venning 2023; Ellison et al. 2018). This process involves a contractile pilus system and an assemblage of competence-accessory proteins (Taton et al. 2020; Ellison et al. 2018; Chen et al. 2020). In cyanobacteria, natural competence is under circadian clock control, with pili biogenesis occurring during the day phase and competence being induced with the onset of the night, coinciding with the peak of the circadian cycle (Taton et al. 2020). In the cyanobacterium *Synechocystis* sp. PCC 6803, the cellular levels of c-di-AMP and the transcription of its synthesizing di-adenylate cyclase gene (*dacA*), also follow a circadian rhythm: They decline during the night and increase sharply at the onset of the day (Selim et al. 2021a).

To test the involvement of c-di-AMP in natural competence, we compared the ability of the wildtype (WT) *Synechocystis* strain and the c-di-AMP-free Δ*dacA* mutant to take up DNA. Both strains were incubated with various extracellular DNA constructs containing a sequence that allows the insertion of a chloramphenicol resistance cassette via homologous recombination in a neutral site in the genome. The transformation efficiency for each strain was determined by analyzing the ability of these cells to grow on agar plates in the presence of chloramphenicol. The Δ*dacA* mutant showed significantly lower transformation efficiency than the WT, implying an essential role of c-di-AMP in natural competence (Figs. 1A and S1). Introducing a plasmid containing the *dacA* gene under the control of the P*petE* promoter into the Δ*dacA* strain restored the transformation efficiency to WT levels (Figs. 1A and S1). Introducing the same plasmid into the WT to generate a c-di-AMP overexpression strain (WT::petE-*dacA*) did not affect the transformation efficiency. These results indicate that the absence of c-di-AMP affects cyanobacterial natural competence negatively, whereas high c-di-AMP does not. A similar result was obtained using the c-di-AMP-null Δ*cdaA* mutant of *Synechococcus elongatus* (Fig. S1) (Rubin et al. 2018), indicating that the c-di-AMP–dependent control of natural competence is a common trait among cyanobacteria.

**Figure 1.**
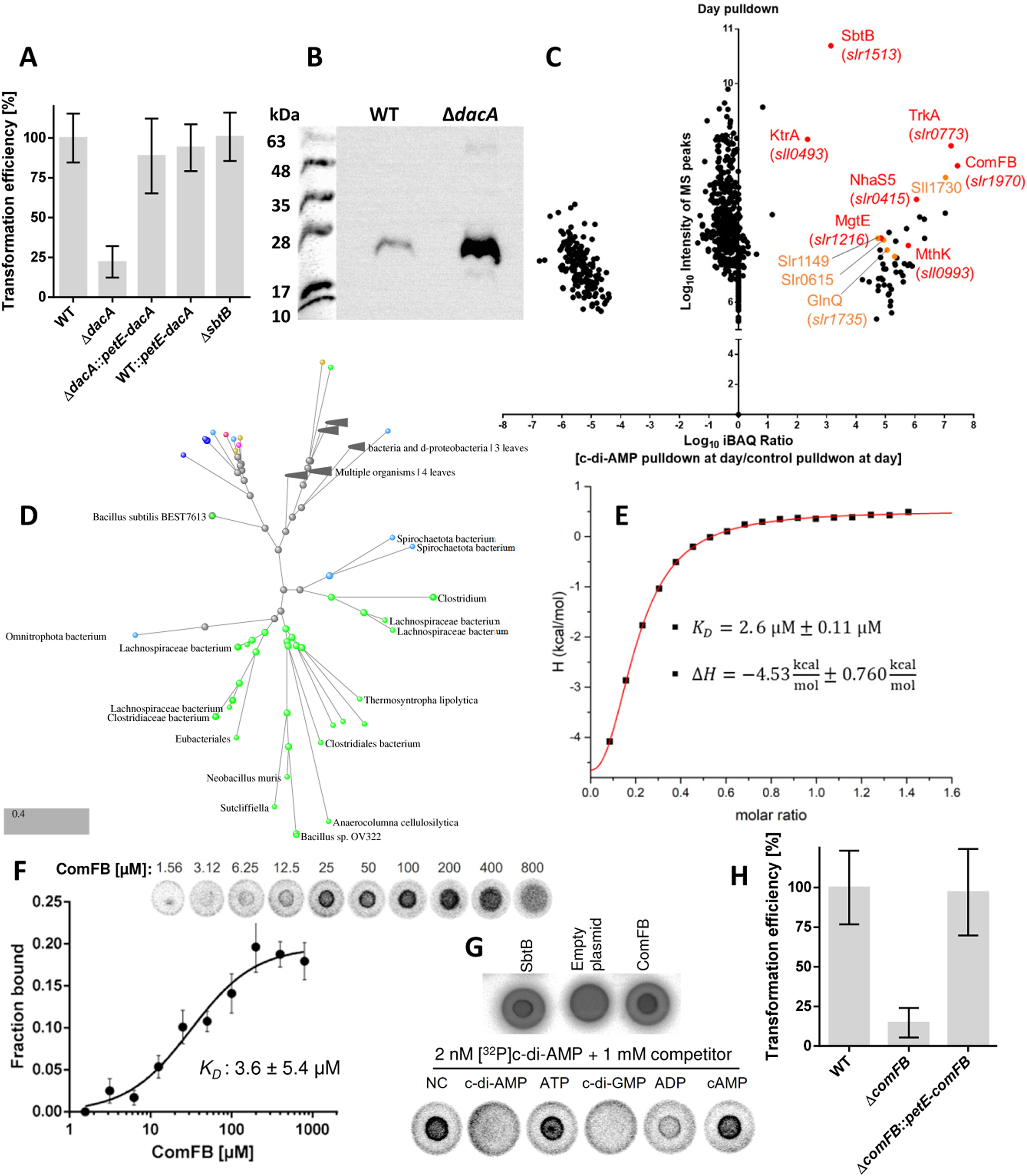
Involvement of c-di-AMP in cyanobacterial natural competence via ComFB signaling protein. **(A)** Transformation efficiency of WT, Δ*dacA*, *dacA::petE-dacA,* WT*::petE-dacA* and Δ*sbtB* strains. **(B)** Immunodetection of PilA1 in the exoproteome of WT and Δ*dacA*. **(C)** Pulldown experiment using immobilized c-di-AMP and extracts of *Synechocystis* cells grown under day-night cycles, showing the enriched proteins in the day phase. Potential new c-di-AMP receptors are highlighted in orange. **(D)** Phylogenetic tree showing the widespread of ComFB proteins among different bacterial phyla (detailed tree see Fig. S4). **(E)** Dissociation constant (*K_D_*) of c-di-AMP binding to ComFB and enthalpy (ΔH) are obtained from sigmoidal fitting curve of all ITC experiments with different monomeric ComFB concentrations (60, 72, 134 and 172 µM). **(F)** DRaCALA assay showing the binding of [^32^P]c-di-AMP to purified ComFB in a concentration dependent manner as indicated. The upper panel shows a representative of one replicate from four technical replicates. The lower panel shows the calculated mean ± SD of the quantification of the bound fraction of [^32^P]c-di-AMP to ComFB from the four replicates and the best fitting curve with the obtained *K_D_* value. **(G)** DRaCALA competition binding assay showing the competition of [^32^P]c-di-AMP with different nucleotides to bind ComFB. NC refers to no competitor. SbtB and cell extract of *E. coli* harboring an empty plasmid were used as positive and negative control, respectively. **(H)** Transformation efficiency of WT, Δ*comFB*, and Δ*comFB::*petE-*comFB* strains.

To gain insights into the molecular basis of c-di-AMP-dependent control of natural competence, we checked how the lack of this second messenger affects pilus biogenesis. A proteome analysis of the Δ*dacA* mutant compared to WT cells under day-night cycle, a condition triggering pili biogenesis and natural competence (Taton et al. 2020), revealed a strong downregulation of specific proteins involved in pilus biogenesis and DNA uptake in the Δ*dacA* mutant (Haffner et al. 2023b). These changes were marked by a reduced abundance of PilT1 (Slr0161), PilM (Slr1274), PilN (Slr1275), PilO (Slr1276), and Sll0180 proteins (Table S1). The cellular levels of the other pilus machinery proteins were not significantly altered in the Δ*dacA* mutant (Table S1). The assembly of a functional pilus requires two motor ATPases, PilB1 and PilT1. PilT1 is located at the pilus base and is required for retraction and depolymerization. Therefore, the *pilT1* mutant is nonmotile, hyperpiliated and loses natural competence (Cengic et al. 2018; Bhaya et al. 2000; Okamoto & Ohmori 2002). The PilMNO proteins form the alignment complex, connecting the components of the pilus machinery in the inner and outer membranes by forming a ring-structure in the periplasm (Chang et al. 2016; Conradi et al. 2020; Chen et al. 2020). Similarly, the *pilM*, *pilN* and *pilO* mutants are nonmotile and non-transformable (Yoshihara et al. 2001). Sll0180 is an accessory protein of TolC-system, needed for the glycosylation of the extracellular pilin PilA1 (Sll1694) and the secretion of the S-layer protein, and thereby the correct assembly of the pilus machinery (Gonçalves et al. 2018). The absence of functional PilA1 causes a non-transformable phenotype (Yoshihara et al. 2001). Notably, our transcriptome analysis showed a partial downregulation of *pilT1* and *sll0180*, while the *pilT2* gene (*sll1533*; Bhaya et al. 2000) was strongly downregulated in Δ*dacA* (Table S2; Mantovani et al. 2022). These findings explain why Δ*dacA* cells lost their natural competence.

The striking decrease of PilT1 levels in Δ*dacA* suggested a strong defect in pilus assembly and retraction. To confirm this assumption, we examined negatively stained Δ*dacA* and WT cells by transmission electron microscopy (TEM). While we could detect both thick and thin pili in the WT, only thick pili were obvious in Δ*dacA* cells (Fig. S2). Additionally, Δ*dacA* mutant showed a hyperpiliation phenotype in analogy to *pilT1* mutant (Cengic et al. 2018; Bhaya et al. 2000; Okamoto & Ohmori 2002). The quantification of the major pilin PilA1 in the exoproteome of the Δ*dacA* mutant via immunodetection revealed an accumulation of PilA1 compared to the WT cells (Fig. 1B), consistent with the hyperpiliation phenotype of Δ*dacA* cells. These findings clearly support the notion of a c-di-AMP-dependent control of pilus biogenesis and natural competence with dysregulation of various components of pilus machinery in Δ*dacA* cells.

The lack of nonretractable pili explains the inability of the Δ*dacA* strain to take up DNA. Interestingly, the downregulation of the above-mentioned proteins was not detected in the Δ*sbtB* mutant, which lacks the only known cyanobacterial c-di-AMP receptor (Table S3) (Selim et al. 2021a; Haffner et al. 2023b). Furthermore, Δ*sbtB* behaved like the WT in our natural competence assays (Figs. 1A and S1). These data suggested the involvement of an additional, yet unknown, c-di-AMP receptor, required for natural competence. To identify this new c-di-AMP receptor, cell extracts of *Synechocystis* cells grown under day-night cycles were incubated with immobilized c-di-AMP and bound proteins were identified by mass spectrometry (Figs. 1C and S3A-C). The identification of several known c-di-AMP targets: SbtB, as well as the transporters TrkA, KrtA, MthK, MgtE and NhaS5, validated our pulldowns (Selim et al. 2021a). Additionally, the Slr1970 protein was also enriched under both conditions. This protein possesses a competence factor B domain (domain PF10719 in the Pfam database) (Mistry et al. 2021) and therefore was annotated as ComFB in the NCBI’s RefSeq database (Haft et al. 2023; Samier et al. 2024). Enrichment of ComFB correlated with the intracellular levels of c-di-AMP, where ComFB was 8 times more abundant in the day than in the night pulldown (Fig. S3C). This indicates that the abundance of ComFB follows the same pattern as the c-di-AMP synthesis with an increase during the day and a decrease at night (Selim et al. 2021a). We found that ComFB proteins are widespread among different bacterial phyla (Figs. 1D and S4), implying a fundamental role in cell physiology (Samir et al. 2024). In *Bacillus, comFB* forms an operon with *comFA* and *comFC*, which are known to be involved in DNA uptake (Samir et al. 2024; Sysoeva et al. 2015; Hahn et al 2019). In *Synechocystis* and other cyanobacteria, *comFB* forms an operon with *hfq,* which is also involved in DNA uptake and motility (Fig. S5) (Dienst et al. 2008; Schuergers et al. 2014; Oeser et al. 2021; Conradi et al. 2020), strongly suggesting a potential function of ComFB in natural competence.

To validate ComFB as a novel c-di-AMP-binding protein, we used several biophysical methods after purifying recombinant His_8_-ComFB protein. Size exclusion chromatography coupled to multiangle light scattering (SEC-MALS) showed a species of ∼ 40.5 kDa (Fig. S6A), indicating that ComFB protein (theoretical mass of the monomer the 20.4 kDa) is dimeric in solution. Using isothermal titration calorimetry (ITC), we measured the binding affinity of ComFB to c-di-AMP. As a control, we also measured ComFB’s binding affinity to ATP, ADP, AMP, cAMP, and cGMP. The c-di-AMP binds endothermically to ComFB with high affinity in the low micromolar range of a K_D_ ∼ 2.6 ± 0.11 μM (Figs. 1E and S6B), while no binding was observed for the other tested nucleotides (Fig. S7), indicating that ComFB binds c-di-AMP specifically. This result was confirmed using nano differential scanning fluorimetry (nanoDSF) and thermal shift assays (Figs. S8 and S9), where c-di-AMP thermally stabilized ComFB significantly in a concentration-dependent manner. Moreover, we performed DRaCALA assays (Roelofs et al. 2011) using radiolabeled [^32^P]c-di-AMP to test further the specificity of c-di-AMP-binding to ComFB in competition with various unlabeled nucleotides. DRaCALA titration assays revealed strong binding of [^32^P]c-di-AMP to ComFB with a K_D_ value of 3.6 ± 5.4 µM (Fig. 1F), comparable to that obtained by ITC. In the competition assays, the unlabeled c-di-AMP competed with [^32^P]c-di-AMP for binding to ComFB, but that was not the case for ATP, ADP, and cAMP, confirming that c-di-AMP binding to ComFB is specific. However, this assay revealed that ComFB could additionally bind c-di-GMP, as c-di-GMP efficiently competed with [^32^P]c-di-AMP (Fig. 1G). Remarkably, a recent study showed that the ComFB homolog in the multicellular cyanobacterium *Nostoc*, named CdgR, controls cell size by binding c-di-GMP (Zeng et al. 2023). In fact, the filamentous cyanobacterium *Nostoc* sp. PCC7120 is regarded as being not naturally competent (Schirmacher et al. 2020), implying that ComFB or CdgR might play different roles in multicellularity lifestyle.

To ascertain whether c-di-AMP binding to CdgR is of physiological relevance, we performed a pulldown assay with immobilized c-di-AMP as described above but using cell extracts from *Nostoc* (Fig. S3D). Indeed, we identified the ComFB homolog (CdgR; Alr3277) as one of the highly enriched proteins along with other known c-di-AMP receptor proteins (Selim et al. 2021a). This result further confirms that both ComFB and CdgR specifically bind both cyclic di-nucleotides in both organisms. Additionally, ComFB was found to bind c-di-GMP with comparable affinity (1.7 ± 0.5 µM) to that of c-di-AMP (Zeng et al. 2023). The existence of a crosstalk between c-di-AMP and c-di-GMP on ComFB awaits, however, further investigation. Crosstalk between second messenger nucleotides is perhaps a more common phenomenon than so far realized. Recently, it was found that the second messengers c-di-GMP and (p)ppGpp reciprocally control *Caulobacter crescentus* growth by competitive binding to a metabolic switch protein, SmbA (Shyp et al. 2021). SbtB plays a similar role in cyanobacterial physiology and binds both cAMP and c-di-AMP (Selim et al. 2018, 2021a, 2023; Selim & Alva 2024; Forchhammer et al. 2022). Likewise, the mycobacterial transcription factor DarR, which is regulated by c-di-AMP-binding, was found to be regulated as well through cAMP-binding (Schumacher et al. 2023). Additionally, crosstalk between cyclic guanosine and adenosine second messengers is also known, as the CRP-Fnr transcription factors are known to bind both cAMP and cGMP, being only active in the cAMP-bound form in *E. coli*, while both cyclic nucleotides mediate the CRP activation in *Sinorhizobium meliloti* (Krol et al. 2023; Werel et al. 2023). Furthermore, both of cAMP and cGMP were found also to bind and modulate the activity of the AphA phosphatase in *E. coli* and *Haemophilus influenzae* (Kronborg & Zhang 2023).

To rule out that the transformation deficiency observed in Δ*dacA* mutant (Fig. 1A) is due to a downstream effect on the intracellular c-di-GMP content, which is known to regulate motility-related functions (Mantovani et al. 2023a; Enomoto et al. 2023), we measured the c-di-GMP levels in Δ*dacA* during the day and the night phases (Fig. S10). The c-di-GMP levels were comparable in both strains within the light/dark phases, thus confirming that DNA uptake is influenced by c-di-AMP specifically.

To clarify if ComFB plays a role in natural competence, we created a Δ*slr1970* deletion mutant (Δ*comFB*; Fig. S11) and tested its DNA natural competence ability as described above (Figs. 1H and S1C). Like Δ*dacA,* Δ*comFB* showed reduced transformation efficiency as compared to the WT and Δ*sbtB* cells. Complementation of Δ*comFB* mutant with *comFB* gene under a P*petE* promoter restored the WT competence phenotype (Fig. 1H). Interestingly, no impairment in DNA uptake was observed for Δ*sbtB*, which lacks another c-di-AMP receptor protein and showed a similar transformation efficiency to the WT cells (Figs. 1A and S1C). These findings further support that natural competence depends on c-di-AMP signaling and is controlled specifically by a pathway that involves ComFB as a c-di-AMP receptor protein, while other c-di-AMP targets are not involved in this process. Notably, in contrast to Δ*sbtB* (Selim et al. 2021a), the Δ*comFB* mutant did not show any impairment under diurnal growth or prolonged dark incubation (Fig. S12), supporting the notion that c-di-AMP plays different signaling functions through binding to different cellular receptors.

To gain insights into the molecular basis on how ComFB controls the natural competence, we carried out a comparative proteome analysis of Δ*comFB* and WT cells, focusing on the motility and the pilus biogenesis-related proteins (Fig. S13), as described above for Δ*dacA*. Surprisingly PilA1 (Sll1694) was the most upregulated protein in Δ*comFB* as compared to the WT, implying a hyperpiliation phenotype in analogy to *dacA* mutant (Figs. S2 and 1B). PilA1 is part of the operon (*sll1693*-*sll1696*) (Sergeyenko & Los 2000; Singh et al. 2005; Linhartová et al. 2021). We were able to detect also upregulation in both of Sll1696 and Sll1693, which is a SAM-dependent methyl transferase potentially involved in methylation in PilA1 and PilA2. Similarly, the minor pilin PilX2 (Slr0442; Oeser et al. 2021) was also upregulated, further supporting the hypothesis of the hyperpiliation phenotype. The S-layer protein (Slr1704) and the outer membrane porin (Sll1550), which are required for the cell envelop and the correct assembly of pilus machinery (Huang et al. 2004; Qiu et al. 2018; Gonçalves et al. 2018), were also upregulated in Δ*comFB*. Moreover, the Deg protease (Sll1679), Sll0141 and Sll1581 were also upregulated in Δ*comFB* mutant. The Deg protease (Sll1679) is known to be involved in motility and piliation (Chen et al. 2020), while the Sll0141 is an accessory protein needed for pilin glycosylation and secretory machinery (Gonçalves et al. 2018) and the Sll1581 is important for the production of cell-surface exopolysaccharides (Jittawuttipoka et al. 2013).

Additionally, PixJ1 (Sll0041) and PixL (Sll0043) of TaxD1, and PilJ (Sll1294) and CheA (Sll1296) of TaxD2, which are motility-related proteins and involved in phototaxis (Bhaya 2004), were down-regulated in Δ*comFB.* In agreement with the upregulation of PilA1, it was reported previously that Δ*pixL* mutant shows upregulation of *pilA1* (Shin et al. 2008). The Δ*pilJ* mutant was shown previously to be non-transformable and non-motile (Yoshihara et al. 2002). Moreover, several proteins (e.g. Sll0445, Sll0446, Slr0362, and Slr0442) which are under the control of LexA, a transcription factor known to regulate motility and pilus biogenesis (Kizawa et al. 2016; Chen et al. 2020), were deregulated in Δ*comFB* mutant. In analogy to Δ*dacA* mutant, we were able also to detect a downregulation in PilN, which is part of *pilMNO* operon, explaining the reduced transformability of both Δ*dacA* and Δ*comFB* mutants. To further confirm this result by another independent method, we analyzed the transcript levels of *pilM* gene using RT-PCR (Fig. S13). In fact, we were able to confirm a downregulation of the *pilM* in both Δ*comFB* and Δ*dacA* mutants (Fig. S14A). We checked also for the expression of *comFB*, *hfq* and *pilB1* genes. As expected, we were not able to detect *comFB* mRNA in Δ*comFB* mutant, while *hfq* showed a normal expression in all strains. Since the abundance of *hfq* mRNA was similar to that of the WT cells, we can safely assume that the *comFB* mutation does not cause a polar effect on the expression of the upstream *hfq* gene, which is required for pilus assembly (Dienst et al. 2008; Schuergers et al. 2014; Conradi et al. 2020), highlighting the specific effect of *comFB* and *dacA* mutations on *pilMNO* operon. As a negative control, we checked for *pilB1*, where we did not detect any changes.

To test whether the Δ*comFB* mutation causes hyperpiliation in analogy to Δ*dacA* mutant, we examined negatively stained Δ*comFB* cells by TEM. TEM analysis of the Δ*comFB* mutant revealed a hyperpiliated cells (Fig. S2), and the quantification of PilA1 in the exoproteome of the Δ*comFB* mutant via immunoblotting revealed a strong accumulation of PilA1 similar to Δ*dacA* cells (Fig. S14B,C). Altogether, these results confirm that c-di-AMP and ComFB are involved in the same pathway, of both being required for pilus biogenesis and natural competence.

Finally, to determine whether the cellular role of ComFB is conserved among cyanobacteria, we created a Δ*comFB* deletion mutant (*Synpcc7942_1924*) in *S. elongatus* (Fig. S11). This strain showed an impaired ability to take up DNA (Fig. S1E,F), confirming a conserved role for ComFB in natural competence. To gain insight into the pathways that ComFB could coordinate in *S. elongatus*, we checked for co-fitness scores of an RB-TnSeq (Random Barcode Transposon Insertion Site Sequencing) library, which indicate likelihood that two genes participate in similar pathways and respond alike under different growth conditions (Wetmore et al. 2015; Price et al. 2018). RB-TnSeq is a high-throughput genetic screening technique used to identify the function of genes by assessing the impact of gene disruptions on cellular fitness under different conditions (Wetmore et al. 2015). The RB-TnSeq library of *S. elongatus* was grown under different stress conditions and the abundance of the mutants in the library is monitored by the barcodes sequencing, allowing to determine the fitness of each mutant under all stress conditions. Genes, which possess co-fitness values >0.75, are considered to possess robust co-fitness and likely to participate in similar pathways (Suban et al. 2024). *S. elongatus* Δ*comFB* mutant showed a very high co-fitness (0.85-0.98) for mainly genes involved in natural competence and pilus machinery (Suban et al. 2022, 2024; Taton et al. 2020), including *pilA, pilA2, pilB1, pilC,* and *rntAB* (*Synpcc7942_2484-2486*) operon (Fig. S15). Similar to *Synechocystis* Δ*comFB*, *S. elongatus* Δ*comFB* showed also a strong association with *pilMNOQ* operon (*Synpcc7942_2450-2453*). Also, several regulatory genes of pilus biogenesis (e.g. *hfq*, *sigF1* and *esbA*) showed strong co-fitness association with Δ*comFB* mutant of *S. elongatus* (Suban et al. 2024). This result further confirms that ComFB is a new player in controlling cyanobacterial natural competence and pilus biogenesis.

In conclusion, our results show that the regulation of pili biogenesis and natural competence is a new unexplored role of c-di-AMP, which requires the receptor protein ComFB. In a broader context, natural competence is a primary mode of horizontal gene transfer, which plays an important role in spreading multidrug resistance. It would therefore be highly interesting to determine whether the influence of c-di-AMP and ComFB signaling in DNA uptake extends to other bacteria, especially those of clinical relevance. Collectively, we identified ComFB as a novel widespread c-di-NMP-receptor protein, which turned out to be a pivotal competence-accessory protein, at least in cyanobacteria, regulating the pili biogenesis.

## Materials and Methods

### Protein production and purification

All the plasmids and primers used in this study are listed in (Table S5). The *slr1970* (encoding ComFB) and *slr1513* (encoding SbtB) from *Synechocystis* sp. PCC 6803 were cloned into the pET-28a vector with Gibson assembly, thereby incorporating a C-terminal His_8_-tag. Positive clones were selected on 50 μg/mL kanamycin agar plates. The expression and purifications of His_8_-ComFB, His_6_-DisA and His_8_-SbtB proteins were achieved as described previously (Selim et al. 2019, 2021b). The proteins were recombinantly produced in *E. coli* strain LEMO21 (DE3) by overnight induction at 20 °C using 0.5 mM IPTG. Cells were lysed by sonication and the soluble proteins (ComFB, DisA or SbtB) were purified by immobilized metal affinity chromatography (IMAC) using Ni^2+^-Sepharose resin (cytiva^TM^), followed by size exclusion chromatography (SEC) using a Superdex 200 Increase 10/300 GL column (GE HealthCare). Protein purity was assessed by Coomassie-stained SDS-PAGE, and protein concentrations were determined using Bradford assay. Analytical SEC coupled to multi-angle light scattering (SEC-MALS) was conducted as described previously to calculate the molar mass of ComFB protein, whereas SbtB was used as a control (Selim et al. 2019, 2020; Walter et al. 2019).

### Synthesis of [^32^P]c-di-AMP

The pGP2563 plasmid (pET19b-based; kindly provided by Jörg Stülke), expressing an active di-adenylate cyclase His_6_-DisA from *Bacillus subtilis* (Mehne et al. 2013), was used to express and purify DisA. Twenty µM or 50 µM DisA were incubated in DisA reaction buffer (40 mM Tris/HCl [pH 7.5], 100 mM NaCl and 10 mM MgCl_2_) with 1 mM ATP at 30 °C overnight with 300 rpm shaking. Samples were centrifuged for 10 min at 14.000 rpm to remove precipitated protein. Then, the supernatant was filtered through Amicon® Ultra – 0.5 mL 10 kDa cutoff centrifugal filters (Merck KgaA; Darmstadt, Germany). To test the enzymatic efficiency of DisA before synthesizing [^32^P]c-di-AMP, DisA was incubated with unlabeled ATP and 15 µl of the reaction product was analysed by thin layer chromatography (TLC; POLYGRAM CEL300 PEI plates) (Macherey-Nagel GmbH & Co. KG, Düren, Germany) using a running buffer of [1 vol. Saturated NH_4_SO_4_ (4.2 M) and 1.5 volumes 1.5 M KH_2_PO_4_ (pH 3.6)] (Fig. S16). Via injecting 5 µl of the reaction product, the activity of DisA was further confirmed by LC-MS (ESI-TOF mass spectrometer; MicrO-TOF II, Bruker) connected to Ultimate 3000 HPLC system (Dionex) on C18 column (Phenomenex, 150×4.6 mm, 110 Å, 5 μm) and using a flow rate of 0.2 mL/min and a 45 min program (for 5 min, 100 % buffer A (0.1 % formic acid with 0.05% ammonium formate), then 30 min of a linear gradient to 40 % buffer B (100 % acetonitrile), and 10 min of column re-equilibration with 100 % buffer A). Data are presented as extracted ion chromatograms (EICs) for ATP and c-di-AMP (Fig. S16). Finally, 250 µCi radiolabeled [^32^P]c-di-AMP was synthesized from [α-³²P]ATP (6000 mCi/µmol) by using His_6_-tagged DisA by Hartmann Analytic GmbH (Braunschweig, Germany).

### *In vitro* protein-ligand binding assays

Binding of recombinantly produced ComFB to c-di-AMP (or other nucleotides: ATP, ADP, AMP, cAMP and cGMP) was analyzed *in vitro* by isothermal titration calorimetry (ITC), thermal shift assay (TSA), and differential scanning fluorimetry (nanoDSF), as described previously (Lapina et al. 2018; Haffner et al. 2023a; Mantovani et al. 2023b). For ITC and TSA, both ComFB and c-di-AMP were dissolved in binding buffer (50 mM Tris/HCl, pH 8.0, 300 mM NaCl, 0.55 mM EDTA). ITC measurements were conducted on a MicroCal PEAQ-ITC instrument (Malvern Panalytical), at 25 °C, with a reference power of 10 μcal/s. Different ComFB protein concentrations in the range of 60-172 μM were titrated against 0.5 or 1 mM c-di-AMP. A control experiment was recorded by titration of c-di-AMP over a cell filled with buffer, to measure the dilution heat of c-di-AMP. Data were analyzed using one set of binding sites model with the MicroCal PEAQ-ITC Analysis Software (Malvern Panalytical) to calculate the dissociation constant K_D_. The dilution heat of the control ITC buffer/c-di-AMP experiments were subtracted from the ComFB/c-di-AMP runs. For reproducibility, different patches of ComFB protein purifications were used in different ITC experiments. TSA measurements were conducted on an iQ5 Real-Time PCR detection system (Bio-Rad). ComFB (10-39 μM) and 0-1.2 mM c-di-AMP were mixed in different ratios, with the addition of 10x SYPRO Orange. All conditions were measured in triplicate in sealed 96-well plates by following the dye’s fluorescence emission over a temperature range of 25 to 99 °C. Data were analyzed with OriginPro software (OriginLab Corporation) and Python. For nanoDSF (Nanotemper), both ComFB and SbtB proteins were diluted in the ITC buffer and used at 1.5 mg/mL concentration with or without 0.5 mM c-di-AMP. The proteins autofluorescence (350/330 nm ratio), as well as light scattering were measured in a temperature range of 30-99 °C to determine the melting curve and the rate of protein unfolding.

### Differential Radial Capillary Action of Ligand Assay (DRaCALA)

The specificity of c-di-AMP binding to ComFB has been verified using DRaCALA (Differential Radial Capillary Action of Ligand Assay) (Roelofs et al. in 2011) using either *E. coli* cell lysate or purified proteins. For cell lysate, ComFB or SbtB has been overexpressed in *E. coli* LEMO21 (DE3), and cell lysates were used for the DRaCALA assays. In the DRaCALA assays, the cell lysates (with total protein concentration of 20 µg) or purified proteins (ComFB or SbtB) were mixed with 2 nM of a radioactively labeled [^32^P]c-di-AMP (∼ 6000 mCi/µmol) for 15 min at room temperature in a binding buffer of (10 mM Tris/HCl [pH 8.0], 100 mM NaCl and 5 mM MgCl_2_). In the competition binding assays, the [^32^P]c-di-AMP was incubated first for 2 mins with the protein or the cell lysates, before adding 1 mM of unlabeled nucleotides (c-di-AMP, c-di-GMP, ATP, ADP, and cAMP) to the reaction mixture for 15 mins. Finally, 10 µL of each mixture were dropped on a nitrocellulose membrane (Amersham^TM^ ProtanTM 0.2 µm NC, Catalogue No10600001, Cytiva Europe GmbH, Freiburg), which binds to the protein while the free ligands diffuse, thereby a radioactive signal appears at the center of the drop application in case of binding of the [^32^P]c-di-AMP to ComFB. After drying, the nitrocellulose membranes were transferred into an X-ray film cassette and the imaging plates (BAS-IP MS 2025, 20 x 25 cm, FUJIFILM Europe GmbH, Düsseldorf, Germany) were placed directly onto the nitrocellulose membrane, then the cassettes were closed and incubated overnight. The next day, the plates were imaged with a Typhoon^TM^ FLA9500 PhosphorImager (GE Healthcare). For [^32^P]c-di-AMP signal quantification, the image analysis software Image Studio Lite Ver 5.2.5 was used. The fraction of bound nucleotide was calculated based on (Roelofs et al. 2011) using the following equation:

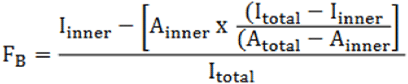

In the DRaCALA competition assays, if one of the unlabeled nucleotides binds to the ComFB, it will compete on the same binding sites and will reduce or eradicate the radioactive signal of [^32^P]c-di-AMP binding to ComFB. SbtB, a known c-di-AMP receptor protein (Selim et al. 2021a), and extract of *E. coli* cells expressing an empty plasmid were used as positive and negative controls, respectively.

### Phylogenetic analysis

Phylogenetic analysis was done essentially as described elsewhere (Neumann et al. 2022). Homologous protein sequences to the ComFB full-length protein (Slr1970, amino acids 1-173) and to the ComFB-domain only (amino acids 57-147) were searched against the Ref-Seq Select proteins database, using the NCBI blastp suite. ComFB hits were filtered (expectation value E ≤ 10^-3^). The sequences were submitted to multiple sequence alignments (MSA) using COBALT (NCBI). Phylogenetic trees were constructed based on the MSAs and visualized using the iTOL online tool (Letunic et al. 2021).

### Construction of mutant strains

The unicellular, freshwater cyanobacterium *Synechocystis* sp. PCC 6803, described in (Selim et al. 2021a), was used as the reference wildtype strain in this study. All plasmids and primers used in this study are listed in (Table S5). All constructs used in this study were generated using Gibson assembly. All knockout mutants were generated with homolog recombination using the natural competence of *Synechocystis* sp. PCC 6803, as described previously (Selim et al. 2018).

For generation of knockout deletion mutants, the mutants were constructed by deleting the ORFs *slr1513*, *sll0505*, and *slr1970* (designated *sbtB*, *dacA*, and *comFB*, respectively) and replaced with the erythromycin, kanamycin, and spectinomycin resistance cassette, respectively. The Δ*sbtB* and Δ*dacA* knockout mutants were created, as described previously (Selim et al. 2018, 2021a). For generation of the knockout mutation in the *slr1970* ORF (designated *comFB*; Fig. S11), a synthetic DNA fragment encoding the upstream and downstream regions of *slr1970* (0.5 kb) and the spectinomycin resistant cassette (gBlock, IDT, USA) were cloned into digested pUC19 vector using the Gibson cloning strategy. For complementation, the Δ*dacA*::petE-*dacA*, WT::petE-*dacA* and Δ*comfB*::petE-*comFB* strains were generated by introducing the *dacA* gene (*sll0505*) or *comFB* gene (*slr1970*) under the control of Cu^2+^ inducible promoter P*petE* into respective mutant backgrounds using the self-replicating plasmid pVZ322, as described previously (Selim et al. 2018, 2021b). All the plasmids used to generate the mutants were verified by sequencing and then transformed in *Synechocystis* sp. PCC6803, as described (Selim et al. 2018). All mutants were selected on BG_11_ plates supplemented with proper antibiotics and verified by PCR.

The c-di-AMP-free mutant in the ORF (*Synpcc7942_0263*) of *Synechococcus elongatus* PCC 7942 was created as described previously (Rubin et al. 2018) using a *cdaA*-deletion plasmid (AM5403; kindly gifted from Susan Golden) carrying the spectinomycin/streptomycin resistance gene *aadA*. For creation of Δ*comFB* (*Synpcc7942_1924*) mutant in *S. elongatus*, 50 mL of *S. elongatus* culture with an OD_750_ of 0.7 were centrifuged at 4200 rpm for 20 min at room temperature. The cell pellet was resuspended in 500 µl of fresh BG11 media. 1 µg of the plasmid pUC19-Δ7942*comFB* was added to the cell suspension. This plasmid contains a spectinomycin resistance cassette flanked by the homologous sequences of *Synpcc7942_1924* that allow recombination at the *comFB* locus. This construct was ordered as a gBlock from IDT (San Diego, USA) and inserted into the pUC19 backbone using Gibson assembly. Cells were incubated with the plasmid in the dark for 24 h and then 250 µl of the cell suspension were spread on a HATF membrane (HATF08250, Sigma-Aldrich, Germany) placed on a BG11-agar plate. After 16 h, the membrane was transferred to a BG11-agar plate supplemented with 15 µg/mL of spectinomycin. After 3 days, the antibiotic concentration was increased to 25 *µ*g/mL, and after additional 3 days to 50 µg/mL. Colonies were visible after 12 days.

### Transformation assay

Natural transformation competence was assessed with different suicide plasmids, all encoding chloramphenicol resistance and integrating into different places in the *Synechocystis* sp. PCC 6803 genome (Oeser et al. 2021). Briefly, the cells of wildtype (WT) *Synechocystis* or respective mutants (Δ*dacA*, Δ*sbtB*, Δ*comFB*, Δ*dacA*::petE-*dacA*, WT::petE-*dacA* and Δ*comFB*::petE-*comFB*) were cultivated in BG_11_ (50 mL) at 28 °C, constant 50 μE m^−2^ s^−1^ and shaking to an OD_750_ of 0.7, then harvested at 4,000 g for 20 min. Cell pellets were resuspended in 600 µl of BG_11_ and all samples were adjusted to the same OD_750_. Cells suspensions were transferred to a 1.5 mL tube and 1 µg of different plasmids, containing chloramphenicol resistance cassette, were added to ensure the reproducibility. The 1.5 mL tubes were covered with aluminum foil and incubated at 28 °C for 3 h, then gently inverted and incubated for 3 more h. 0.45 µm HATF membranes (HATF08250, Sigma-Aldrich, Germany) were placed on BG_11_ plates and 200 µl of the cell suspensions were spread on them. Plates were incubated at 28 °C and 50 μE m^-2^ s^−1^ for 16 h and membranes were then transferred to BG_11_ plates supplemented with 15 µg/mL of chloramphenicol. After 48 h of incubation at 28 °C and 50 μE m^-2^ s^−1^, membranes were transferred to BG_11_ plates supplemented with 30 µg/mL of chloramphenicol and further incubated until singles colonies were visible. At least three-five biological replicates were used for each strain. Some clones were verified by PCR for the insertion of chloramphenicol cassette into the genome. The natural transformation competence for *S. elongatus* strains (WT, Δ*cdaA* and Δ*comFB* mutants) was done as described for *Synechocystis* but using a plasmid carrying kanamycin resistance cassette.

### Exoproteome analysis

*Synechocystis* sp. PCC 6803 wildtype, Δ*dacA* and Δ*comFB* cells were grown in 250 mL of BG_11_ at 28 °C, constant 50 μE m^−2^ s^−1^ and shaking to an OD_750_ of 0.8. Cultures were spun down at 4,000 g for 20 min and the supernatant was filtered through cellulose nitrate membrane filters (7182-004, Cytiva, Marlborough, MA, USA) and concentrated to 1 mL using Amicon Ultra-15 centrifugal filters with a cutoff of 10 kDa (UFC901024, Sigma-Aldrich). Three biological replicates were prepared for each strain. Immunoblot detection of PilA1 in the exoproteome extracts was done as previously described (Oeser et al. 2021) using α-PilA1 antibody (kindly provided by Roman Sobotka; Linhartová et al. 2014).

### Transmission electron microscopy (TEM)

Cells (WT, Δ*dacA* and Δ*comFB*), growing BG_11_ at 28℃ under continuous illumination 50 μE m^−2^ s^−1^, were negatively stained with 2% aqueous uranyl acetate (w/v). Imaging was done with Hitachi HT7800 operated at 100 kV, equipped with an EMSIS Xarosa 20-megapixel CMOS camera (Oeser et al. 2021). Acquired images were analyzed with ImageJ.

### Mass spectrometry-based proteomics analysis

The full proteomics analysis of the *Synechocystis* sp. PCC 6803 wildtype, Δ*sbtB*, Δ*dacA* and Δ*comFB* cells, growing under day-night cycles, was done as described previously in (Haffner et al. 2023b). The full proteomics data sets of Δ*sbtB* and Δ*dacA* mutants are described in (Haffner et al. 2023b). The full proteomic data sets of Δ*comFB* mutant is described in this manuscript in details (Table S4). For the wildtype (WT) strain and Δ*comFB* mutant, the proteome analysis of three independent replicates were performed and displayed high reproducibility based on protein abundance correlations (Fig. S17). More than 1730 proteins could be identified in our proteome dataset (Table S4). The overall protein abundances in the Δ*comFB* mutant was slightly divergent from those of WT cells, as indicated by PCA (Fig. S18), indicating profound changes in the proteome as a result of Δ*comFB* mutation.

The pulldown experiments to identify the potential c-di-AMP target proteins was done as described previously (Selim et al. 2021a) using *Synechocystis* sp. (under day and night conditions) and *Nostoc*. sp. PCC 7120 cell extracts. The determination of intracellular c-di-GMP concentration in wildtype and Δ*dacA* cells was done as described in (Selim et al. 2021a) using mass spectrometry calibrated with ^13^C ^15^N -c-di-GMP and ^13^C ^15^N -c-di-AMP (200 ng/ml each).

## Data availability

The mass spectrometry proteomics data have been deposited to the ProteomeXchange Consortium via the PRIDE partner repository with the dataset identifier PXD045008.

## Acknowledgements

The project was funded by grants from the German Research Foundation (DFG) as part of the priority research programs (SPP1879) and (SPP2389; SE 3449/1-1) to KAS and MH, and by the collaborative research center SFB1381 (project number: 403222702) to KAS, SAV and SSiv. KAS gratefully acknowledges Karl Forchhammer for continued support, and acknowledges the infrastructural support by the Cluster of Excellence “Controlling Microbes to Fight Infections (CMFI)” (EXC2124–390838134) and by the Excellence Strategy of the German Federal and Baden-Württemberg State Governments (Projektförderung: PRO-SELIM-2022-14). We would like to thank Jörg Stülke (Göttingen University), Annegret Wilde (Freiburg University), Michael Galperin (NCBI), Susan Golden (University of California San Diego) and Roman Sobotka (South Bohemia University) for valuable comments and/or sharing materials with us, and the staff of the isotope laboratory and the proteome center (Tübingen University) for their excellent support. We would like to thank also the EM facility at the Faculty of Biology, University of Freiburg, for access to the TEM. The TEM (Hitachi HT7800) was funded by the DFG grant (project number: 426849454) and is operated by University of Freiburg (Faculty of Biology) as a partner unit within the Microscopy and Image Analysis Platform (MIAP) and the Life Imaging Center (LIC). We are also grateful to Heinz Grenzendorf, Markus Burkhardt, Filipp Oesterhelt and Marina Borisova (Tübingen University), and Reinhard Albrecht (MPI for Biology Tübingen) for the excellent support and/or assistance.

## Author contributions

KAS conceived, initiated, designed and supervised the research; SSam, SD, EZ, MH, LD, TM, SPL, SSiv and KAS performed research; SVA supervised the TEM analysis; SSam, SD, EZ, and KAS analyzed data and prepared the figures; and SD and KAS wrote the manuscript. All authors approved the final version of the manuscript.

---

**Fig. S1:**
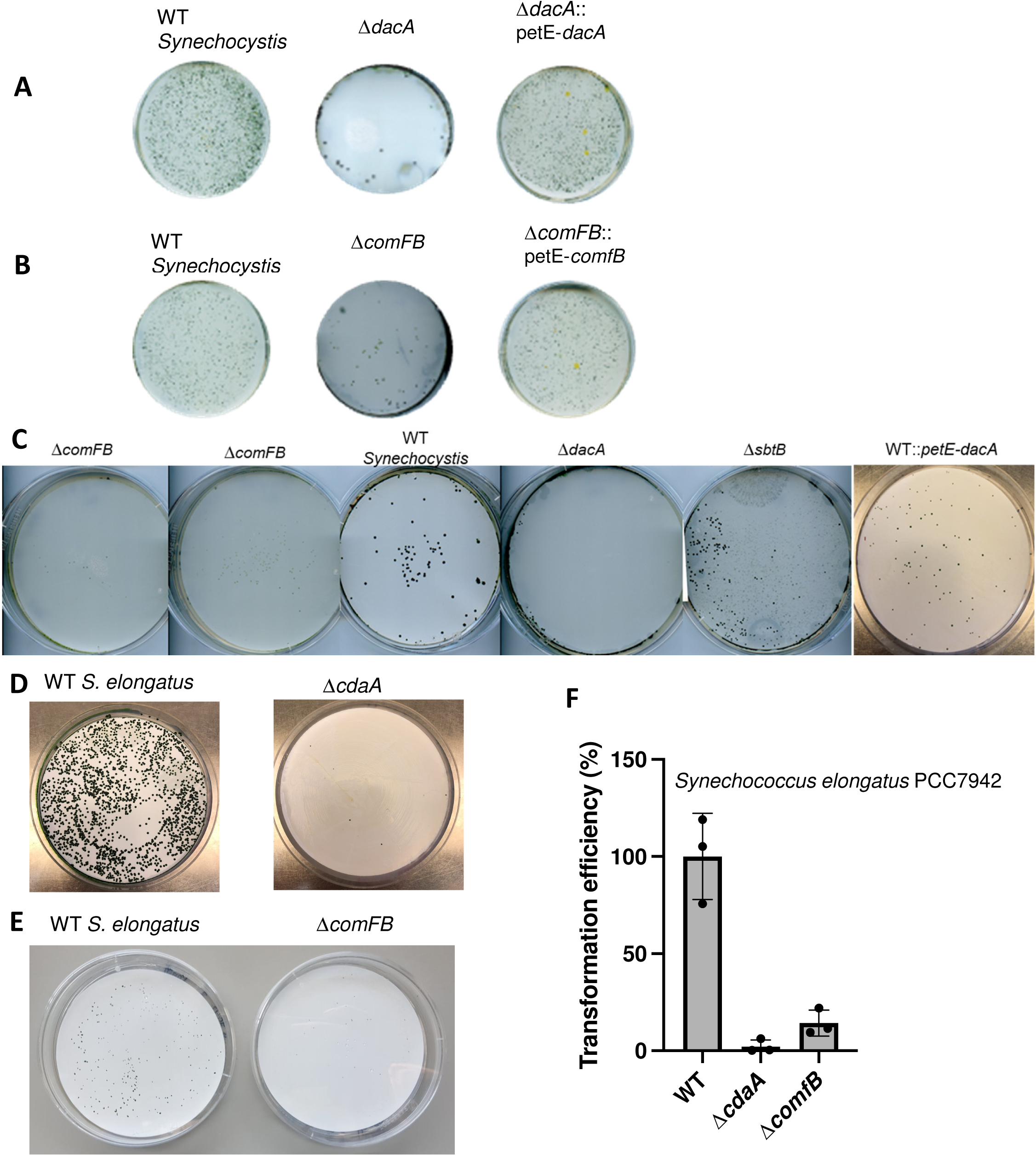
Transformation efficiency of Δ*comFB* and c-di-AMP-free mutants in *Synechocystis* and *Synechococcus elongatus strains*. (A-C) Representative of transformation efficiency in colony forming unites of *Synechocystis* wildtype (WT), Δ*dacA*, *dacA::*petE-*dacA,* Δ*comFB*, and *comFB::*petE-*comFB* strains using different plasmids with chloramphenicol resistance cassette. Δ*sbtB* mutant was used as a negative control, representing another c-di-AMP receptor protein. (D,E) Representative of transformation efficiency in colony forming unites of *S. elongatus* WT and the mutants of Δ*cdaA* (D) and *ΔcomFB* (E). (F) Transformation efficiency of *S. elongatus* strains (WT, Δ*cdaA and ΔcomFB*).

**Fig. S2:**
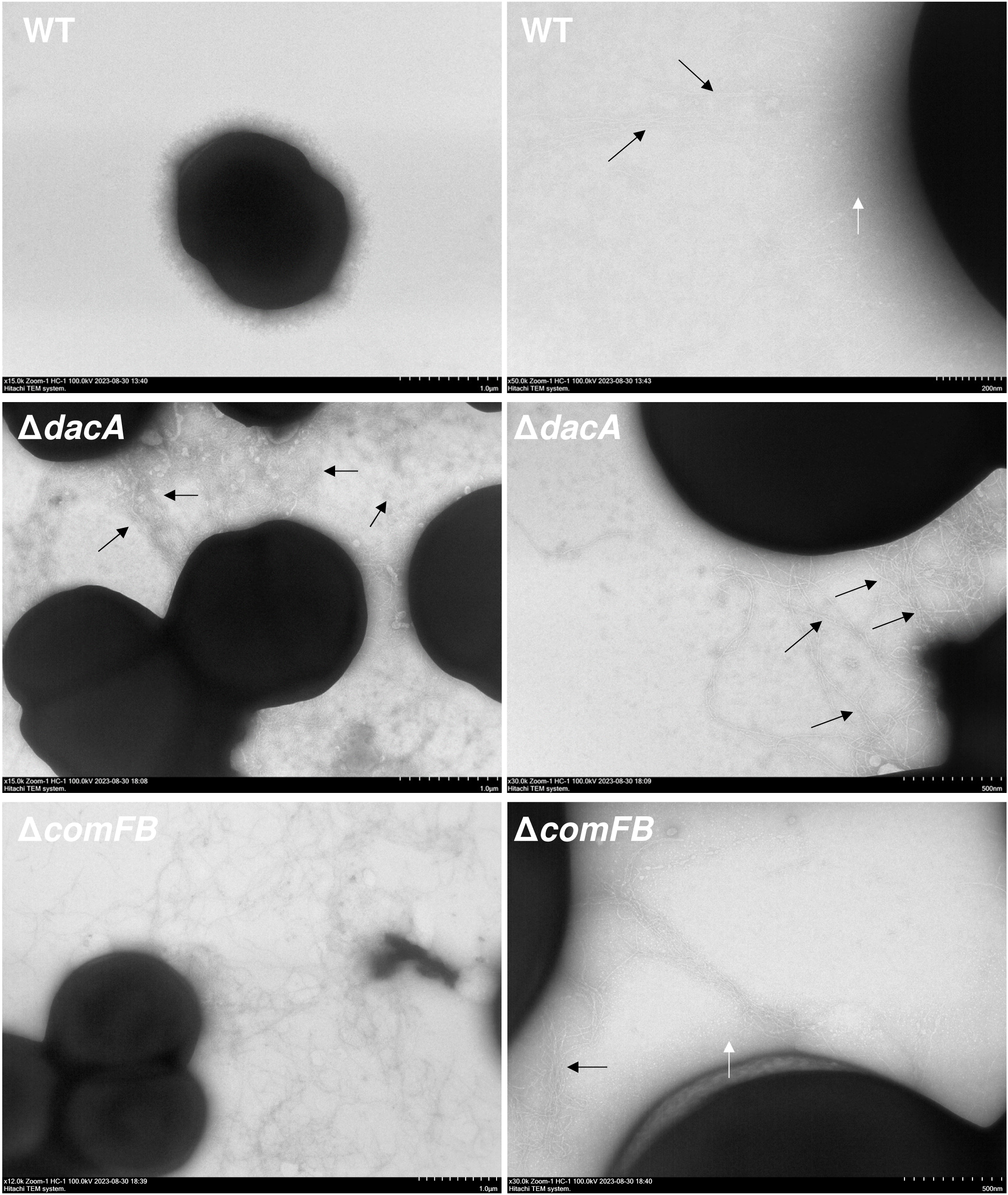
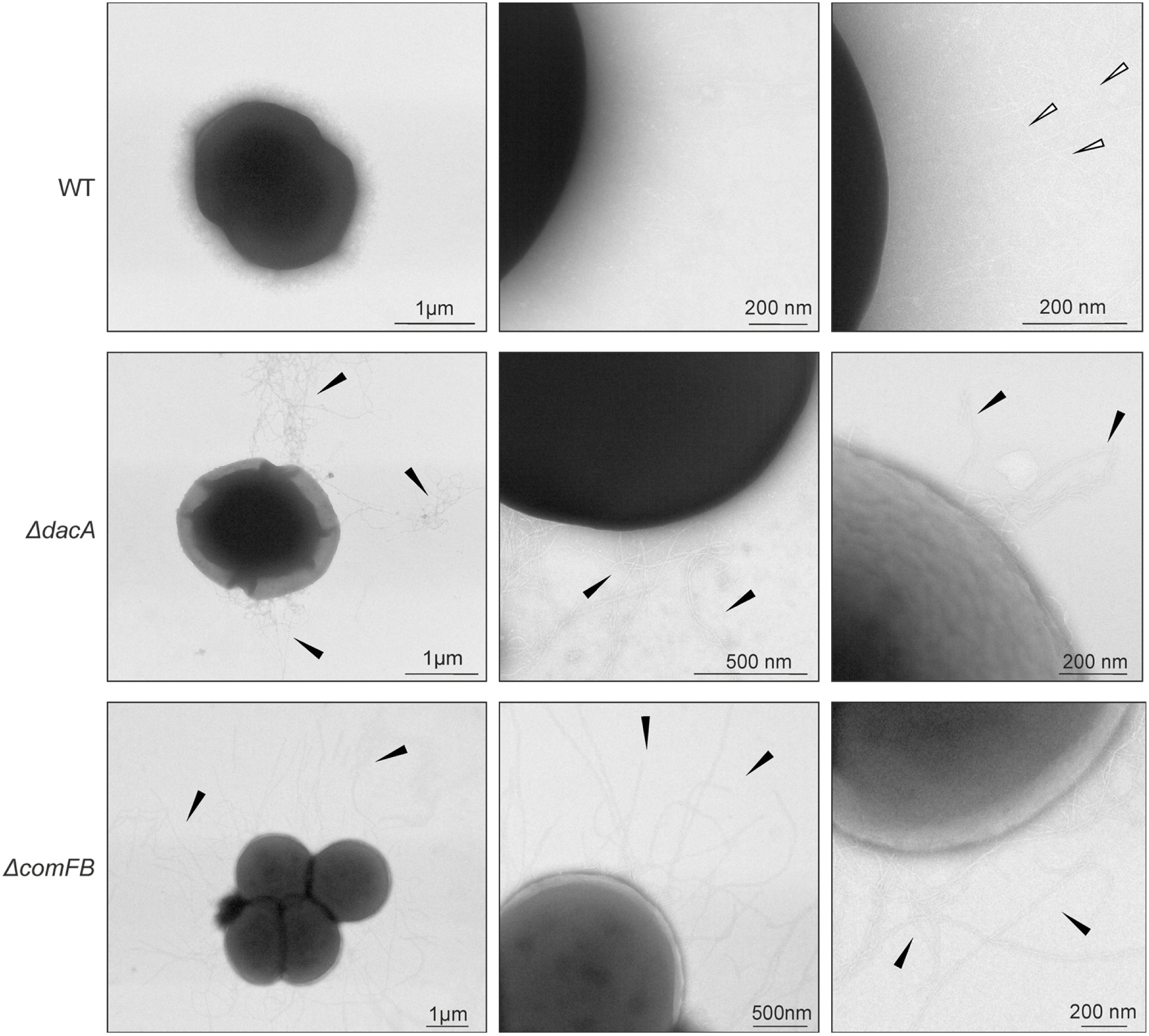
Electron micrographs of negatively stained *Synechocystis* wildtype (WT), Δ*dacA* and *ΔcomFB* cells. Whole cells are depicted with 1 µm scale bar and ultrastructural details of pili are shown in 200-500 nm with distinct types of thick pili (black arrow) and thin pili (white arrow). See also addatinal examples in Fig. S2 (*continue*). **Another example of electron micrographs of negatively stained *Synechocystis* wildtype (WT), Δ*dacA* and *ΔcomFB* cells.** Whole cells are depicted with 1 µm scale bar and ultrastructural details of pili are shown in 200-500 nm with distinct types of thick pili (black arrow) and thin pili (white arrow).

**Fig. S3:**
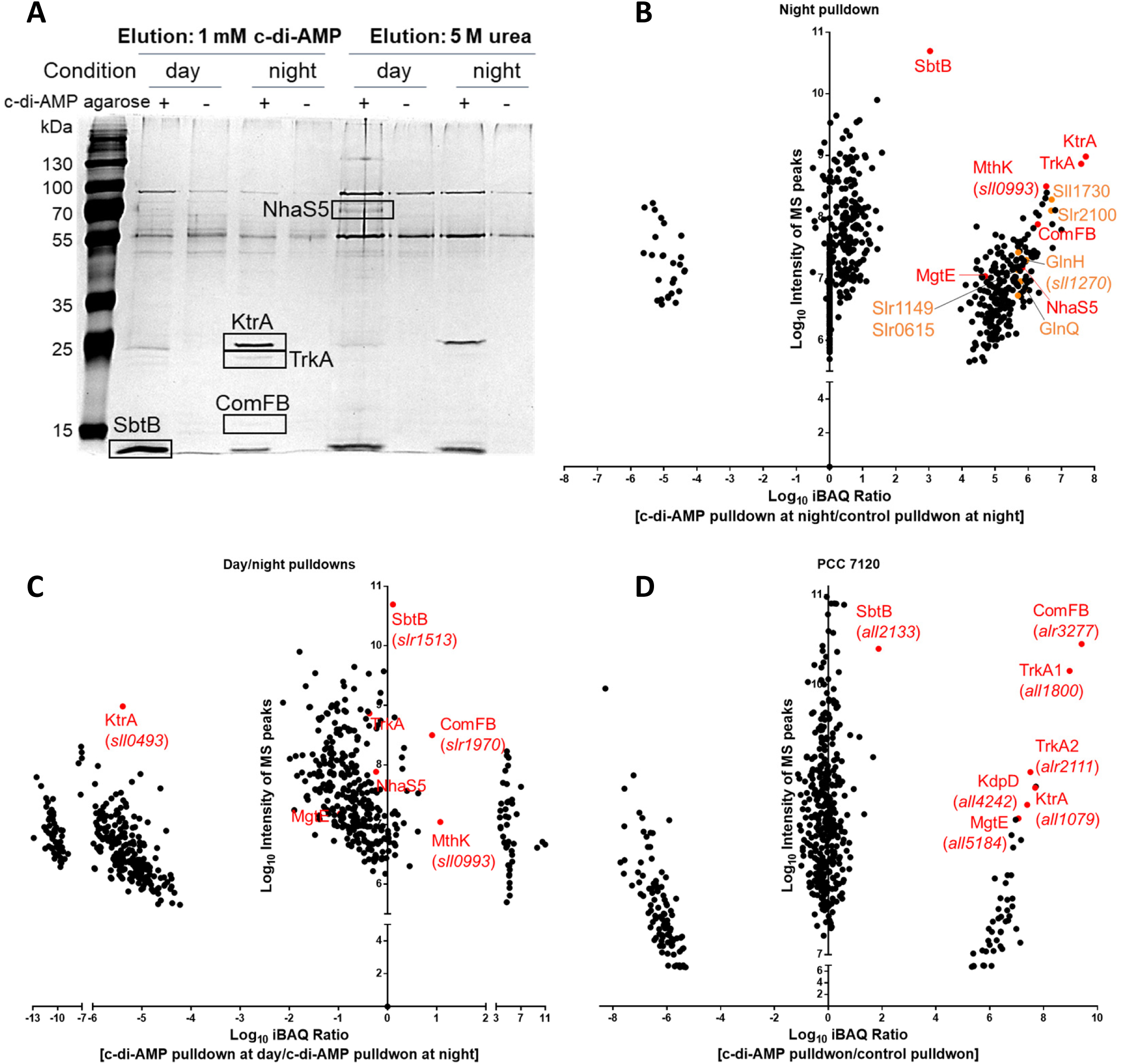
Identification of potential c-di-AMP binding proteins in *Synechocystis* sp. PCC 6803 and *Nostoc* sp. PCC 7120 using immobilized c-di-AMP pulldowns and analyzed by MS-proteomics. (A) SDS-PAGE of c-di-AMP pulldown elution fractions in *Synechocystis* sp. PCC 6803 under day and night conditions, as indicated, with highlight of the potential targets. Elution of bound proteins was achieved by using 1 mM c-di-AMP or 5 mM urea. (B) Identification of potential c-di-AMP binding proteins in *Synechocystis* sp. PCC 6803 under night, enriched proteins are highlighted. (C) enrichment of ComFB in day pulldown compared to night pulldown. (D) Identification of potential c-di-AMP binding proteins in *Nostoc* sp. PCC 7120, enriched proteins are highlighted in red. (B-D) Eluates were analyzed by high accuracy LC-MS/MS to calculate protein enrichment ratios. The identified proteins were sorted by score and refined manually to remove unspecific binning proteins. Significantly enriched proteins were calculated based on Log10 of iBAQ ratio and plotted against the intensity of MS peaks of the identified/defined peptides. The known c-di-AMP receptors: SbtB, TrkA, MgtE and KtrC validated our pulldown approach in general. Potential new c-di-AMP receptors are highlighted in orange.

**Fig. S4:**
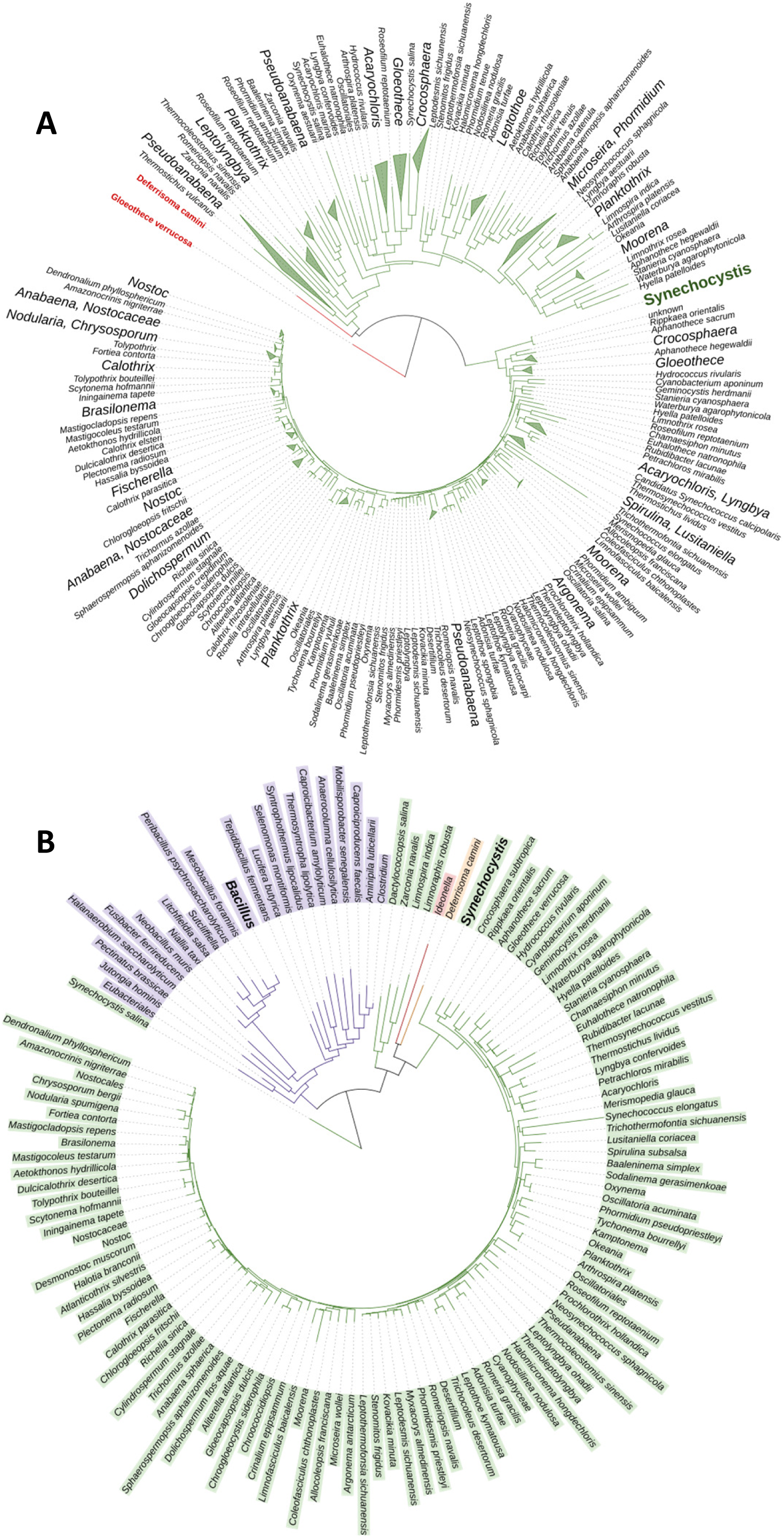
Rooted phylogenetic tree of ComFB homologs. (A) Distribution among cyanobacteria (green). Collapsed clades are shown as triangles. (B) Distribution among cyanobacteria (green), firmicutes (violet), b-proteobacteria (red) as well as other bacteria (orange). Branch lengths represent genetic divergence. Constructed based on a multiple sequence alignment with sequences obtained from a blastp search of the ComFB domain of Slr1970.

**Fig. S5:**
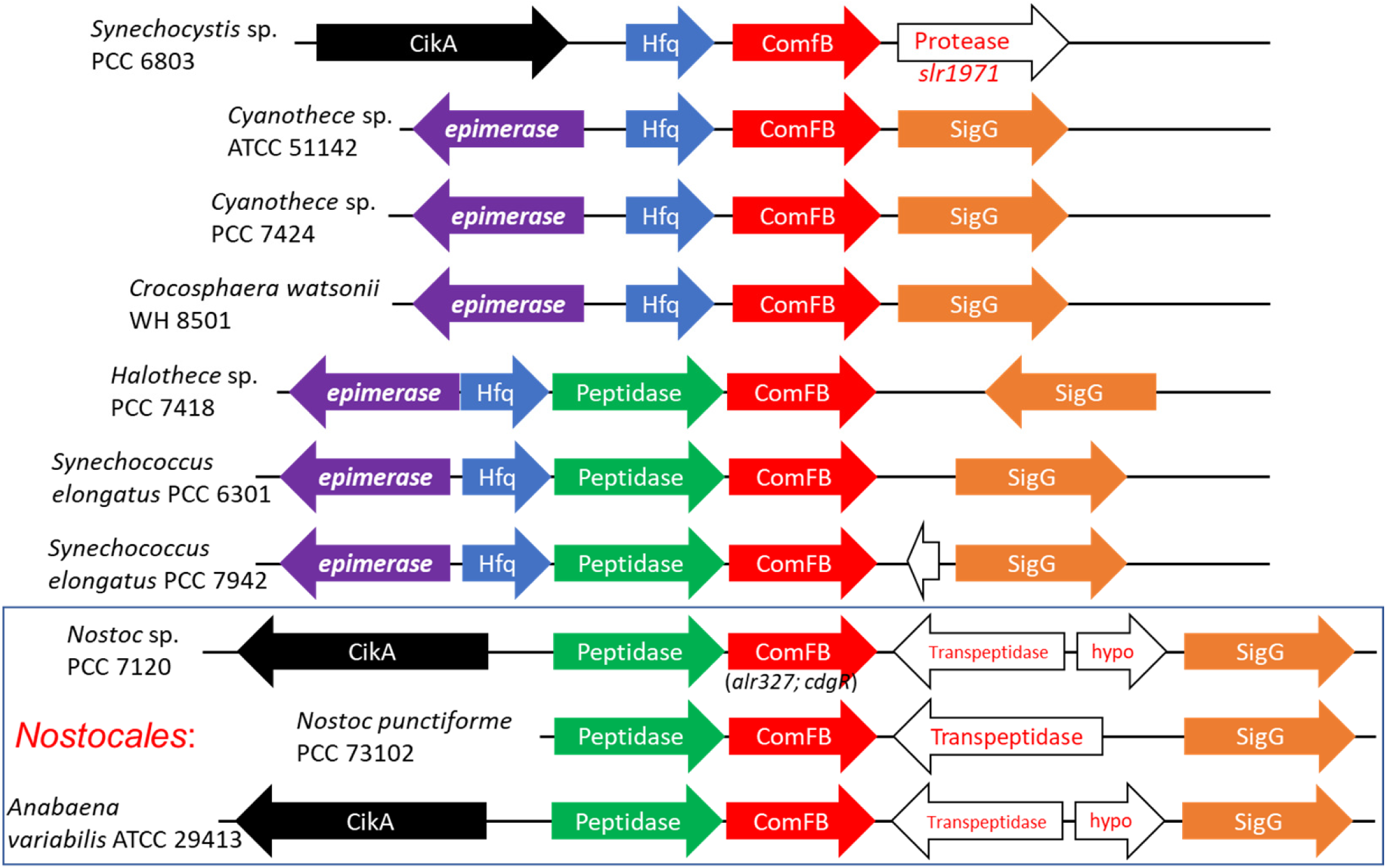
Genomic organisation and conservation of *comFB* homologs (in red) using SEED database in different cyanobacteria species, as indicated. In *Synechocystis* sp. PCC 6803, upstream of *comfB* (*slr1970*; in red) the open reading frames of *hfq* (*ssr3341*; in blue) and *cikA* (*slr1969;* in black) are found. In other cyanobacterial species, the ORFs (open reading frames) encoding for the RNA polymerase sigma factor (SigG; in orange), putative peptidase (in green) and diaminopimelate epimerase (in violet) are found in association with *comfB* as well. Further ORFs which show no strong conservation (e.g. L,D-transpeptidase) or of hypothetical proteins (hypo) are coloured white. The ORF coding for an orthologue of the RNA chaperone Hfq (*ssr3341*) is found to be conserved upstream of ComFB homologs in the unicellular cyanobacterial species, while it seems absent from the multicellular filamentous cyanobacteria of order *Nostocales*. Hfq protein is essential for phototaxis and natural competence, which depends on type IV pili (Dienst et al. 2008). Hfq regulates these processes by binding to the PilB1 ATPase subunit of pili machinery (Schuergers et al. 2014). Another conserved ORF found in association with *comFB* is CikA (*slr1969;* circadian input kinase A), encoding for a photoreceptor regulator of the circadian clock in cyanobacteria (Cohen & Golden 2015; Narikawa et al. 2008). The sigma factors are also involved in circadian clock regulation (Nair et al. 2002). Since ComFB is found in genomic organization with Hfq and circadian clock components, it seems logical that these proteins are also related in their function. Therefore, this further implies the involvement of ComFB in the regulation of light-dependent processes like natural competence or phototaxis (Taton et al. 2020; Menon et al. 2021), which are type IV pili dependent processes.

**Fig. S6:**
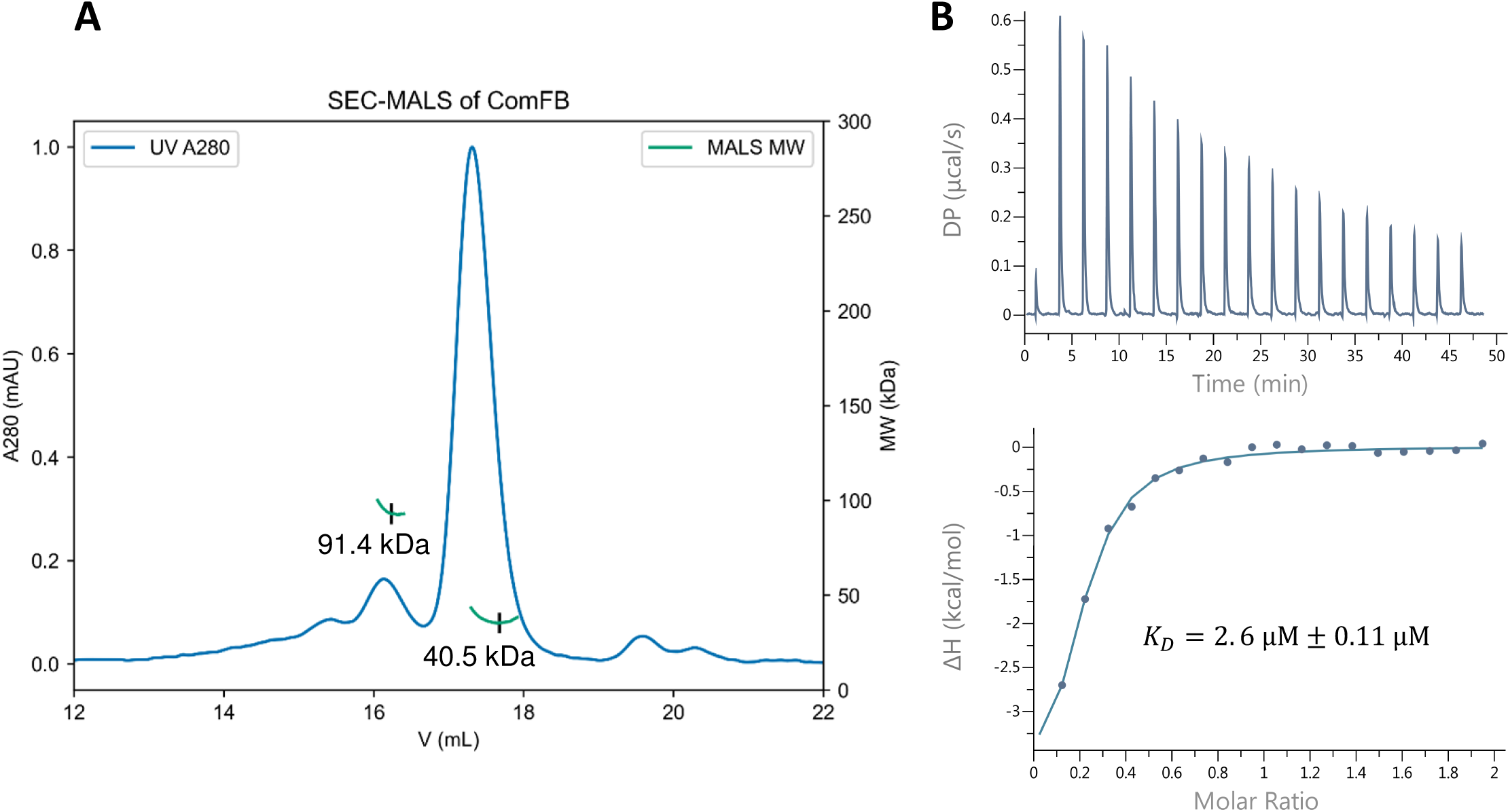
Characterization of ComFB protein encoded by *slr1970*. (A) Size exclusion chromatography coupled to multiangle light scattering (SEC-MALS) of recombinantly purified ComFB. Absorption at 280 nm (A280) is plotted against elution volume (V) from a Superose 6 Increase 10/300 GL column. Molecular weight (MW) obtained from MALS is plotted for the two main peaks in the A280 signal, with minima in the MW marked. The major peak of ComFB (thermotical mass of monomer 20.6 kDa) showed a ∼ 40.5 kDa molar mass, indicating that ComFB behaves as a dimmer in solution, however a small fraction of the protein behaved as a tetramer as indicated by 91. kDa molar mass. (B) Representative isothermal titration calorimetry (ITC) measurements of 172 μM ComFB (monomeric concentration) titrated with 1 mM c-di-AMP. Upper panel shows the recorded differential power (DP) signal of ligand-to-protein titration, plotted against time. The enthalpy changes for each injected (ΔH) are calculated by subtraction of a differential power signal from buffer-in-protein titration control, and subsequent integration of the DP peaks, and plotted against molar ratio of ligand to protein. Lower panel shows the binding isotherms and the best-fit curves according to the one-set of binding sites for dimeric ComFB with K_D_ of 2.6 ± 0.11 μM.

**Fig. S7:**
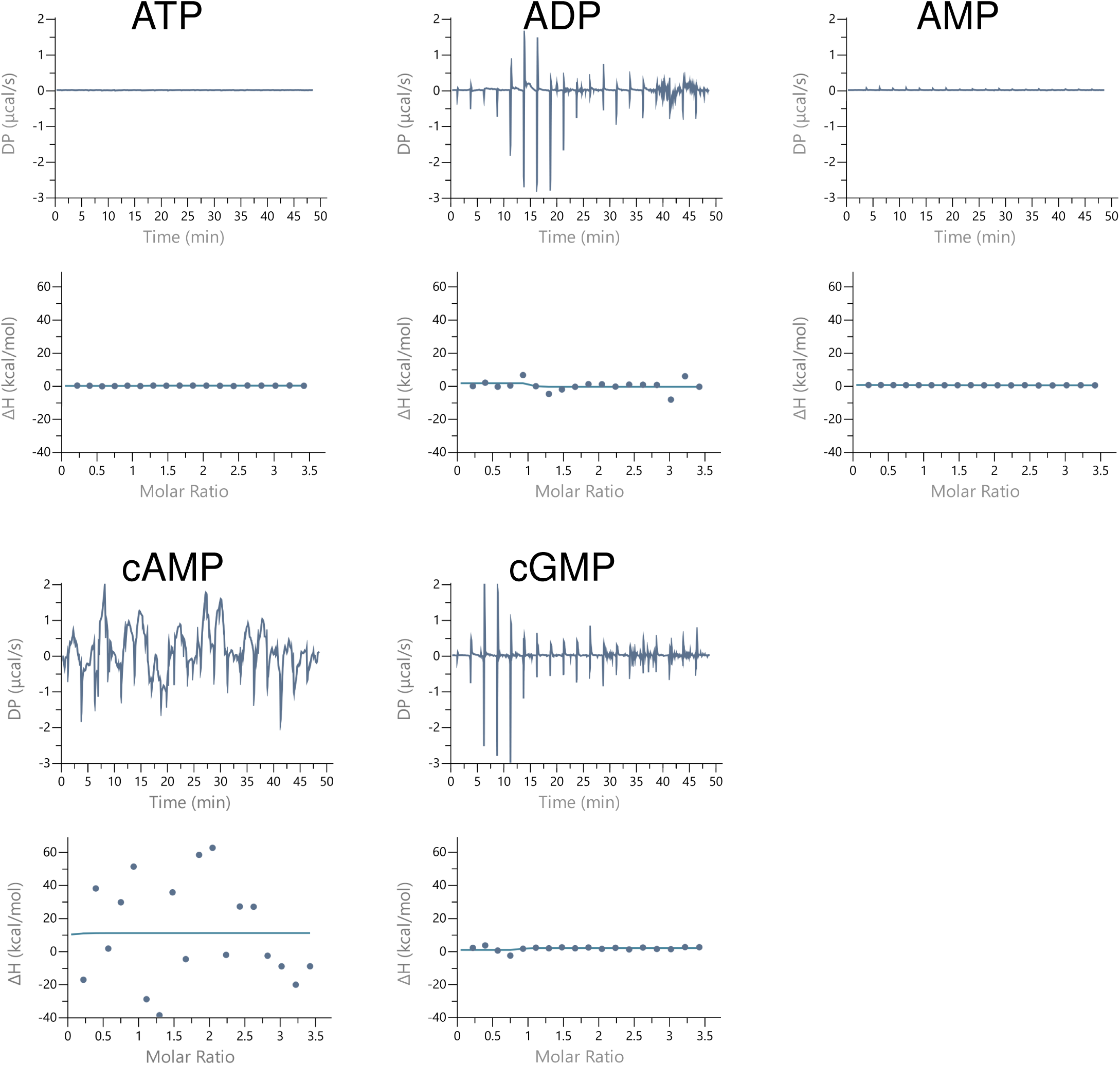
Binding analysis of ComFB to different nucleotides. Representative isothermal titration calorimetry (ITC) measurements of 114 μM ComFB (monomeric concentration) titrated with 1 mM of different nucleotides, as indicated. The Upper panel shows the recorded differential power (DP) signal of ligand-to-protein titration, plotted against time. The lower panel shows the binding isotherms and the fit curves according to the one-set of binding sites for ComFB. All of the tested ligands showed no binding to ComFB, confirming the specificity of c-di-NMP binding to ComFB.

**Fig. S8:**
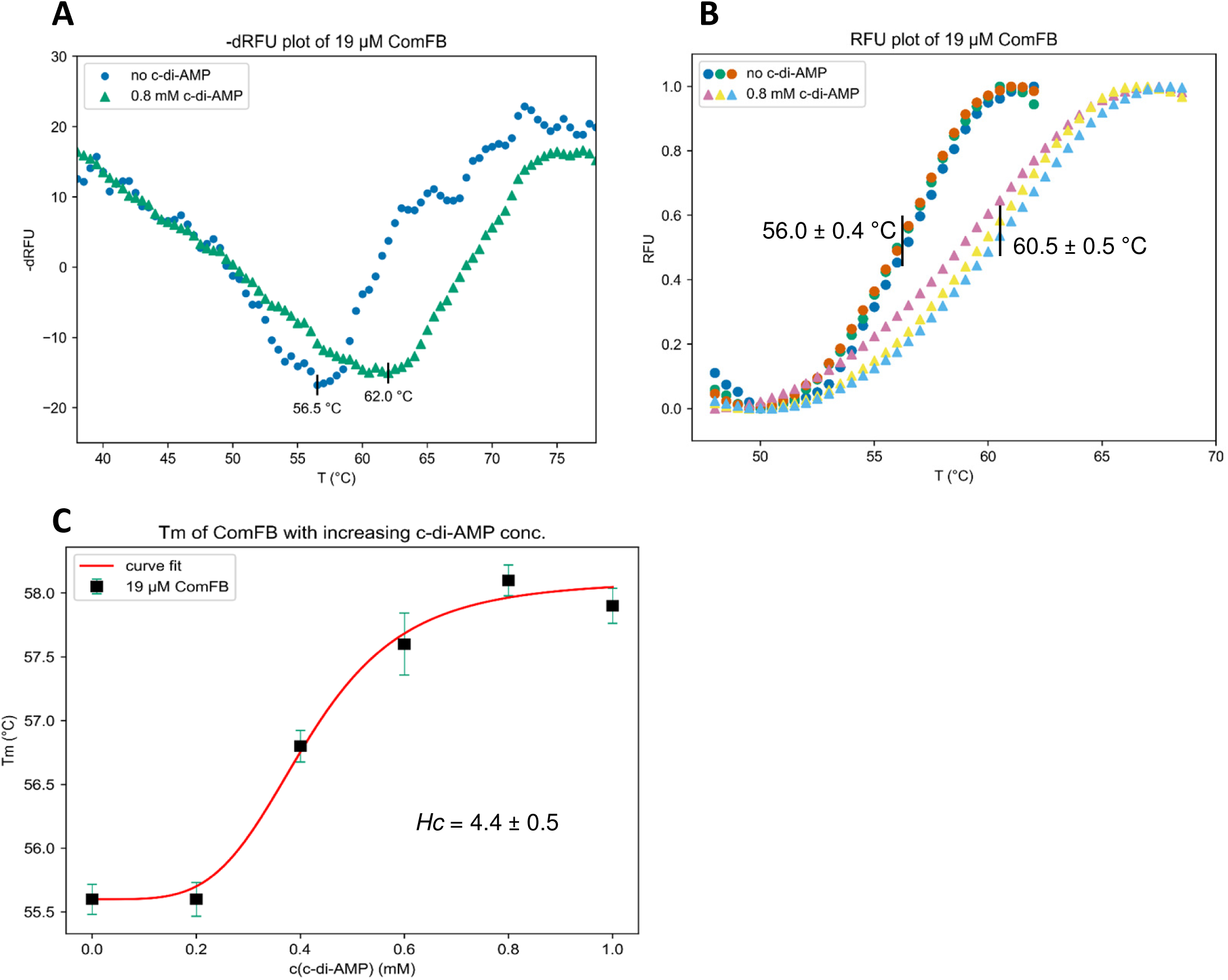
Thermal shift assay showing binding of c-di-AMP to ComFB. (A) Negative first derivative of representative ComFB (19 μM) melting profiles with and without c-di-AMP (0.8 mM), calculated from thermal shift assay data. Minima in the calculated negative first derivative of recorded relative fluorescence units (-dRFU) over temperature (T) represent the melting temperatures of the protein. (B) Melting profile of 19 μM ComFB with and without 0.8 mM c-di-AMP, recorded in a thermal shift assay. The fluorescence emission of SYPRO Orange at 570 nm was followed over a temperature range of 25-99 °C, and normalised relative fluorescence units (RFU) were plotted against temperature (T). Temperatures at half-maximal normalised fluorescence emission are indicated. Measurements were performed in triplicates. (C) Melting temperatures (Tm) of 19 μM ComFB in the presence of different concentrations of c-di-AMP; Melting temperatures were calculated from -dRFU/dT plots as in (A). Measurements were performed in triplicates, error bars show the standard deviation from the calculated mean melting temperature. The data were fitted with the Hill equation. Hill coefficient (*Hc*) is in positive value, indicative of cooperativity between binding sites. Calculated values of the fitting parameters and their variance are indicated.

**Fig. S9:**
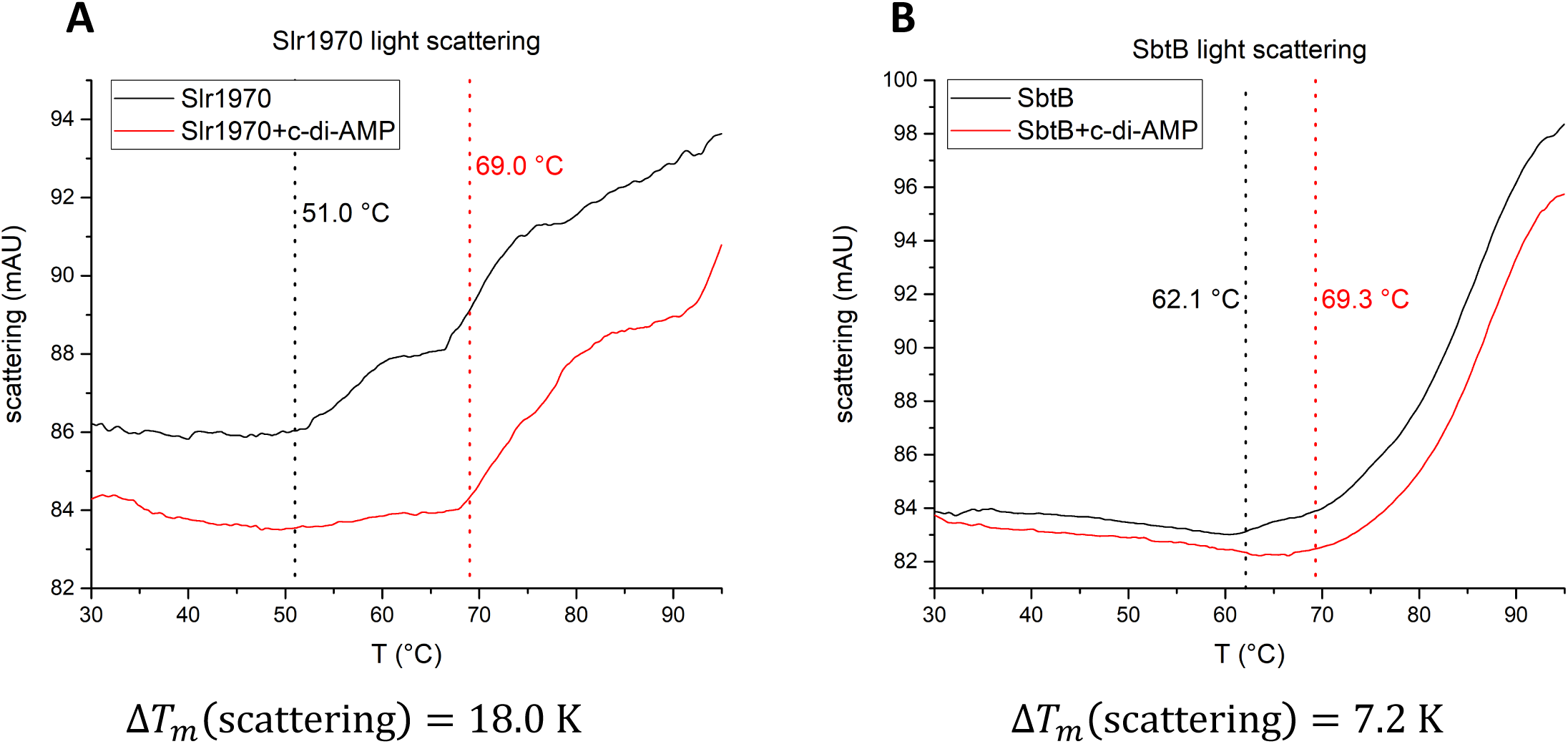
Light scattering obtained from the thermal shift assay using nanoDSF. The calculated temperatures from which the light scattering increases are shown as indicated. The temperature shift (ΔT_m_) between proteins (1.5 mg/ml) with and without c-di-AMP (0.5 mM) is shown as indicated. (A) Light scattering of ComFB (Slr1970), while (B) light scattering of SbtB (used as +ve control as known c-di-AMP receptor protein) (Selim et al. 2021).

**Fig. S10:**
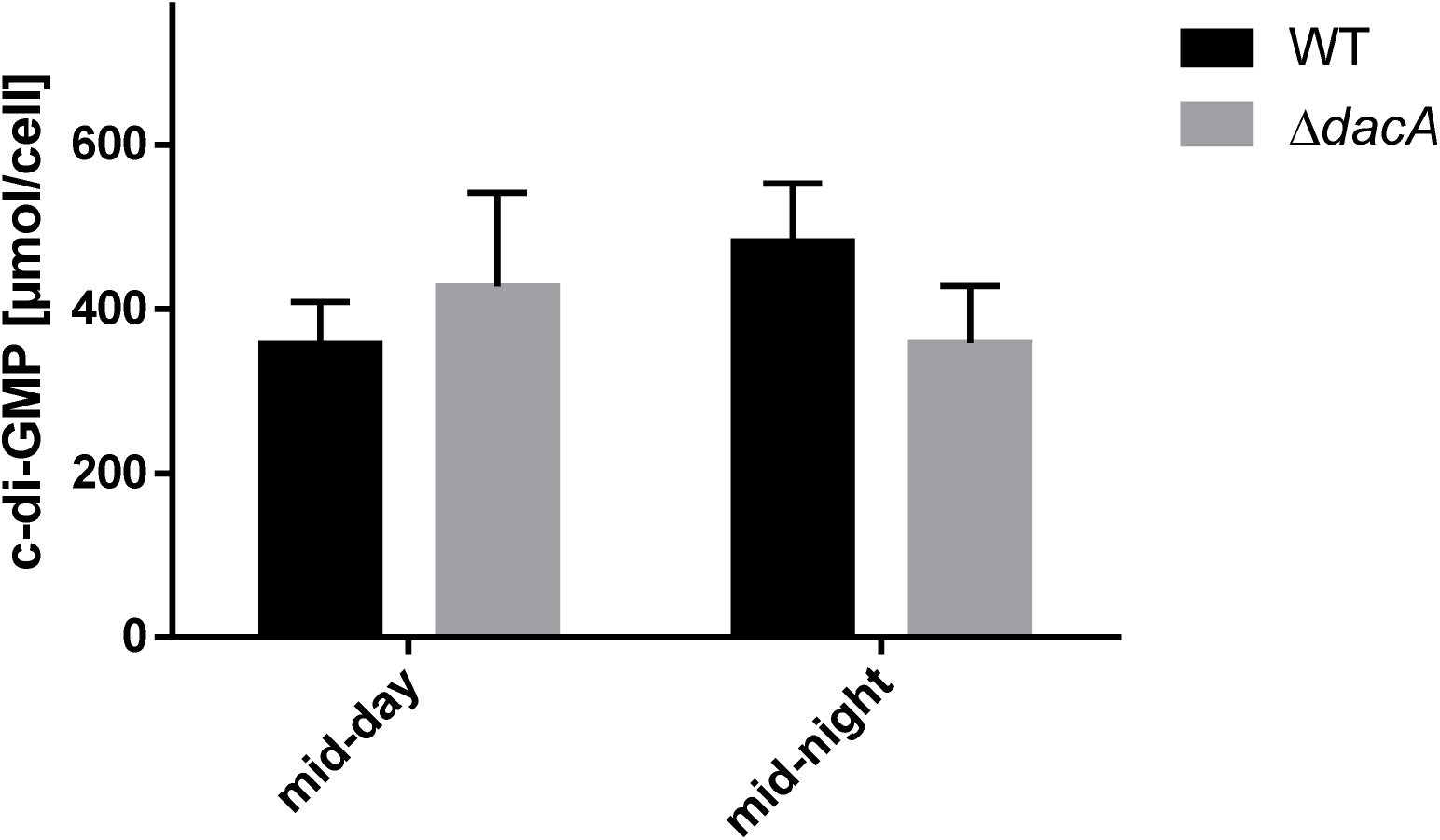
c-di-GMP concentration throughout a 12 h diurnal rhythm within *Synechocystis* WT (black bars) and ΔdacA cells (gray bars) at either mid of day or night phase (i.e. 6 h of light or darkness). X-axis shows the time in hours; Y-axis shows the intracellular concentrations of c-di-GMP.

**Fig. S11:**
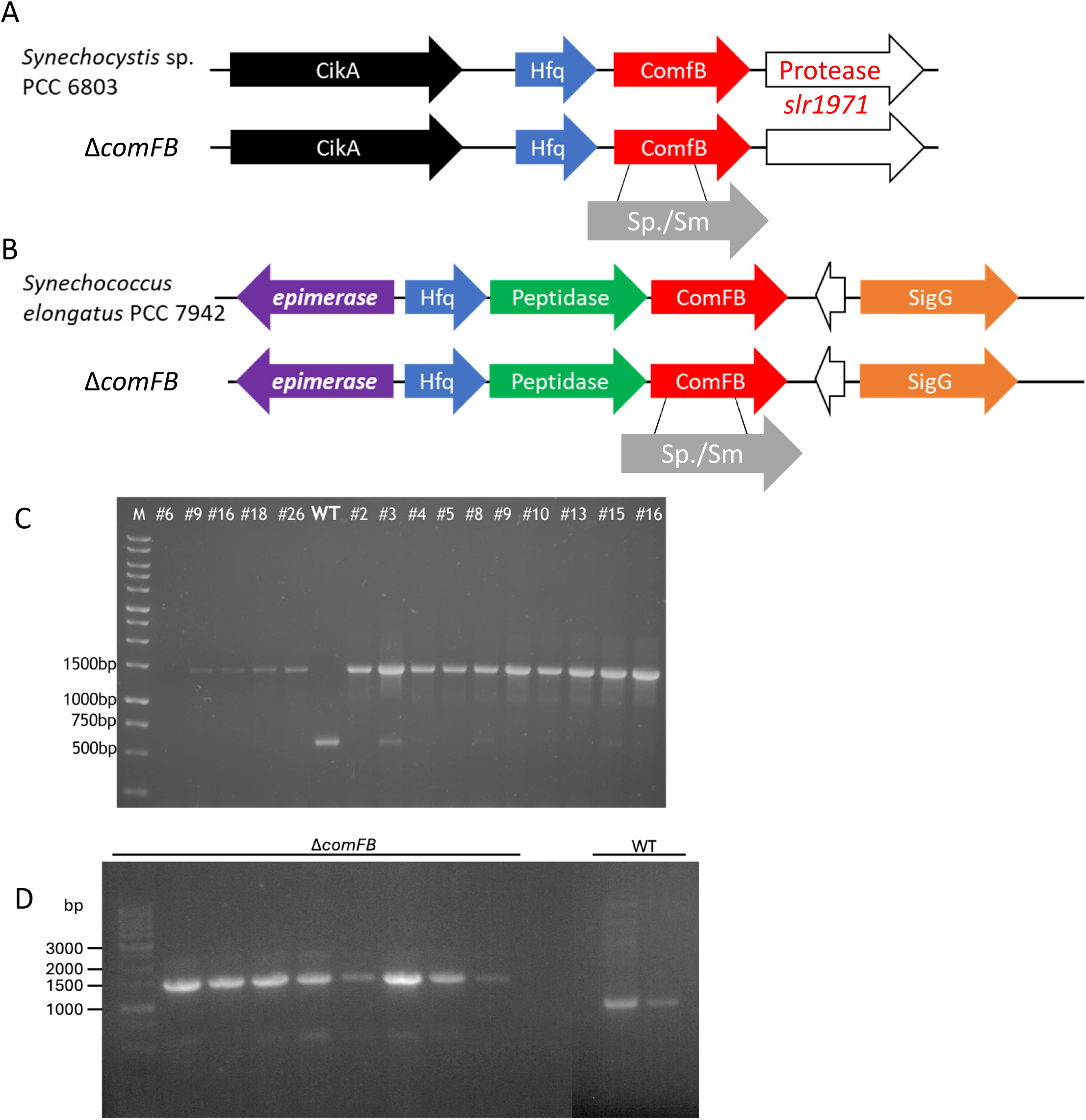
Genotypic characterization of Δ*comFB* knockout mutant in *Synechocystis* sp. PCC 6803 and *Synechococcus elongatus* PCC 7942. (A,B) Schematic representation of genetic organization of *slr1970* and *Synpcc7942_1924* (designated *comFB*) genes in *Synechocystis* sp. PCC 6803 (A) and *S. elongatus* PCC 7942 (B) genomes, respectively. The deletions of the *comFB* gene is achieved by a replacement with spectinomycin/streptomycin (Sp./Sm.) resistance cassette. (C,D) PCR showing complete segregation of Sp./Sm-resistance-cassette in the Δ*comFB* knockout mutants in *Synechocystis* (C) and *S. elongatus* (D) from independent colonies. The PCR product for the Δ*comF*B knockout mutants are around 1500 bp and appears higher than the corresponding WT band, indicating the insertion of Sp./Sm. resistance cassette.

**Fig. S12:**
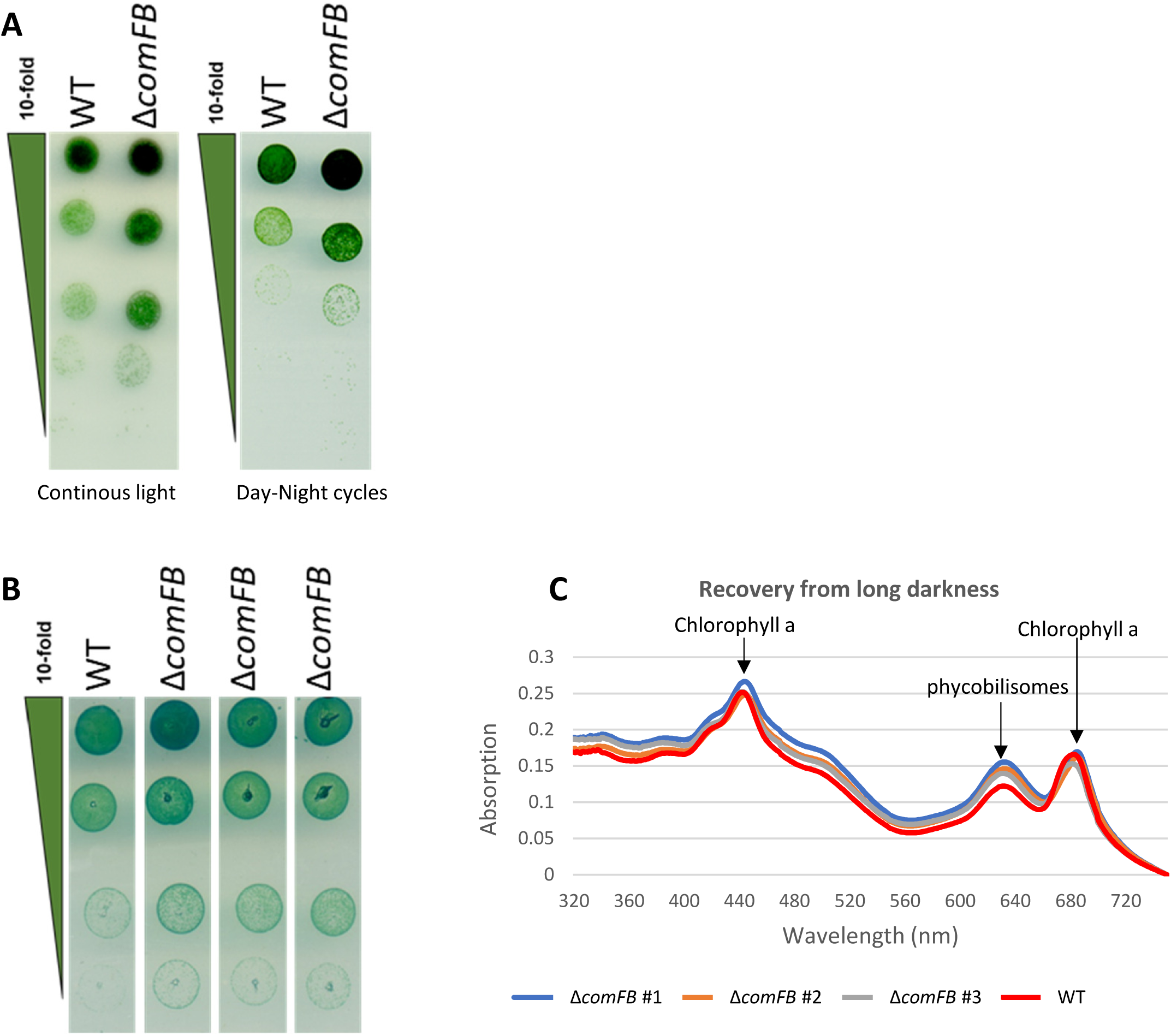
Phenotypic characterization of Δ*comFB* under different light conditions. (A) Growth test by drop plate assay of *Synechocystis* WT and Δ*comFB* cells under either continuous light (left) or a 12-hour diurnal rhythm (right). (B) Viability test using the drop-plate assay of *Synechocystis* WT and Δ*comFB* (3 independent clones of the mutant) cells after 6 days of incubation in complete darkness. Cells were normalized to an optical density at 750 nm (OD_750_) of 1.0 and serial diluted in 10-fold steps (top to bottom; depicted by a green triangle). (C) Whole cell spectra of *Synechocystis* WT cells in comparison to Δ*comFB* cells after 2 days of recovery from darkness (6 days). The peak representing phycobilisomes as well as the peaks representing chlorophyll a are depicted by black arrows. Cultures were normalized to similar OD_750_.

**Fig. S13:**
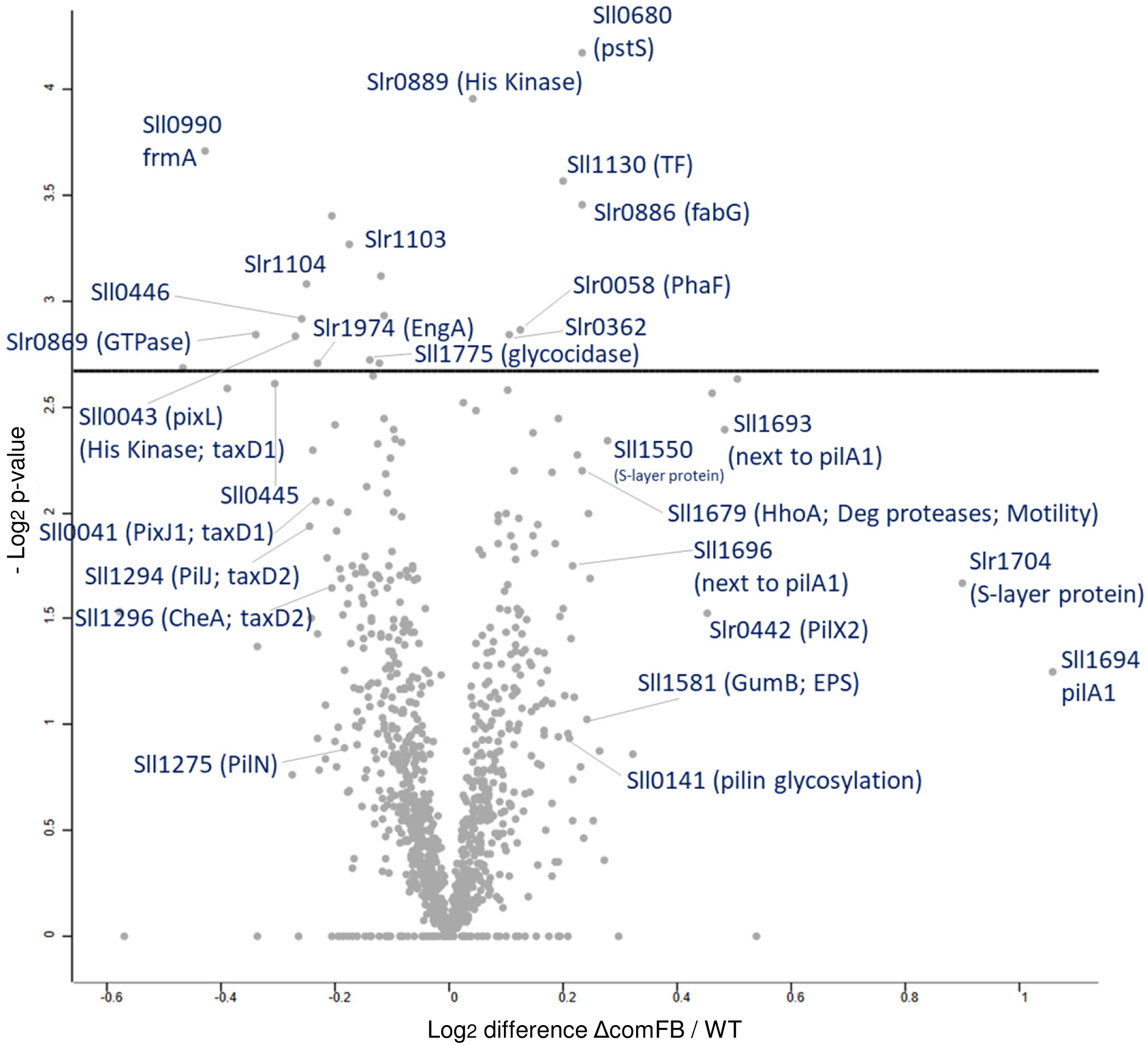
Proteome alterations of Δ*comFB* mutant compared to *Synechocystis* wildtype (WT) cells. Quantitative comparison of protein abundance between Δ*comFB* and WT with corresponding volcano plots indicate differences in protein abundance (Log_2_ difference Δ*comFB*/WT) and corresponding −log_10_ p-values from *t*-test of three independent replicates per strain. Proteins with significant changes in abundance are labeled with positive or negative values, indicating upregulation or downregulation in Δ*comFB* mutant, respectively.

**Fig. S14:**
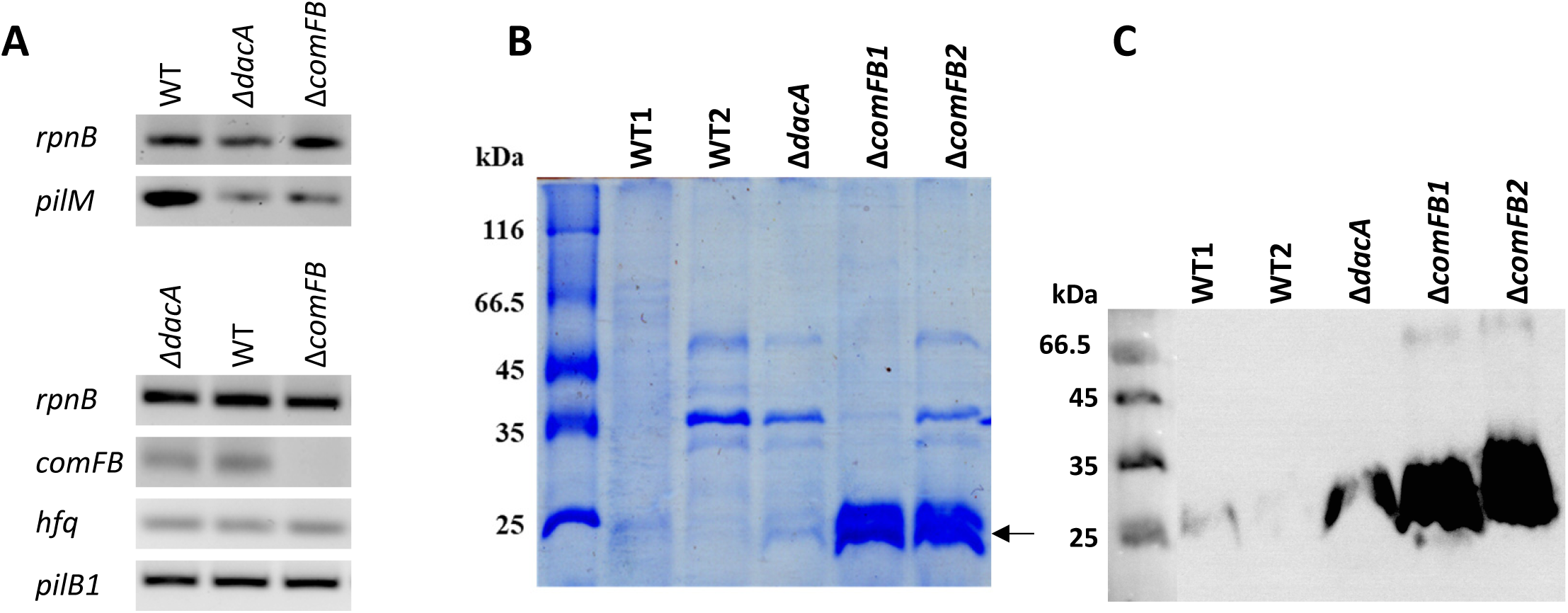
Characterization of *Synechocystis* Δ*dacA* and Δ*comFB* mutants. (A) Gene expression of selected pilus machinery genes in *Synechocystis* wildtype (WT) and Δ*dacA* and Δ*comFB* mutants analyzed using semiquantitative RT-PCR. The constitutively expressed *rnpB* gene served as loading control; a representative of 2 independent biological replicates is shown. (B) Exoproteome of WT, Δ*dacA* and Δ*comFB* cells, separated on SDS-PAGE and stained by coomassie blue and showing accumulation of PilA1 (indicated by black arrow). (C) Immunodetection of PilA1 in the exoproteome of WT and Δ*comFB* mutant for 2 independent biological replicates (indicated by 1 and 2). The Δ*dacA* mutant was used as positive control.

**Fig. S15:**
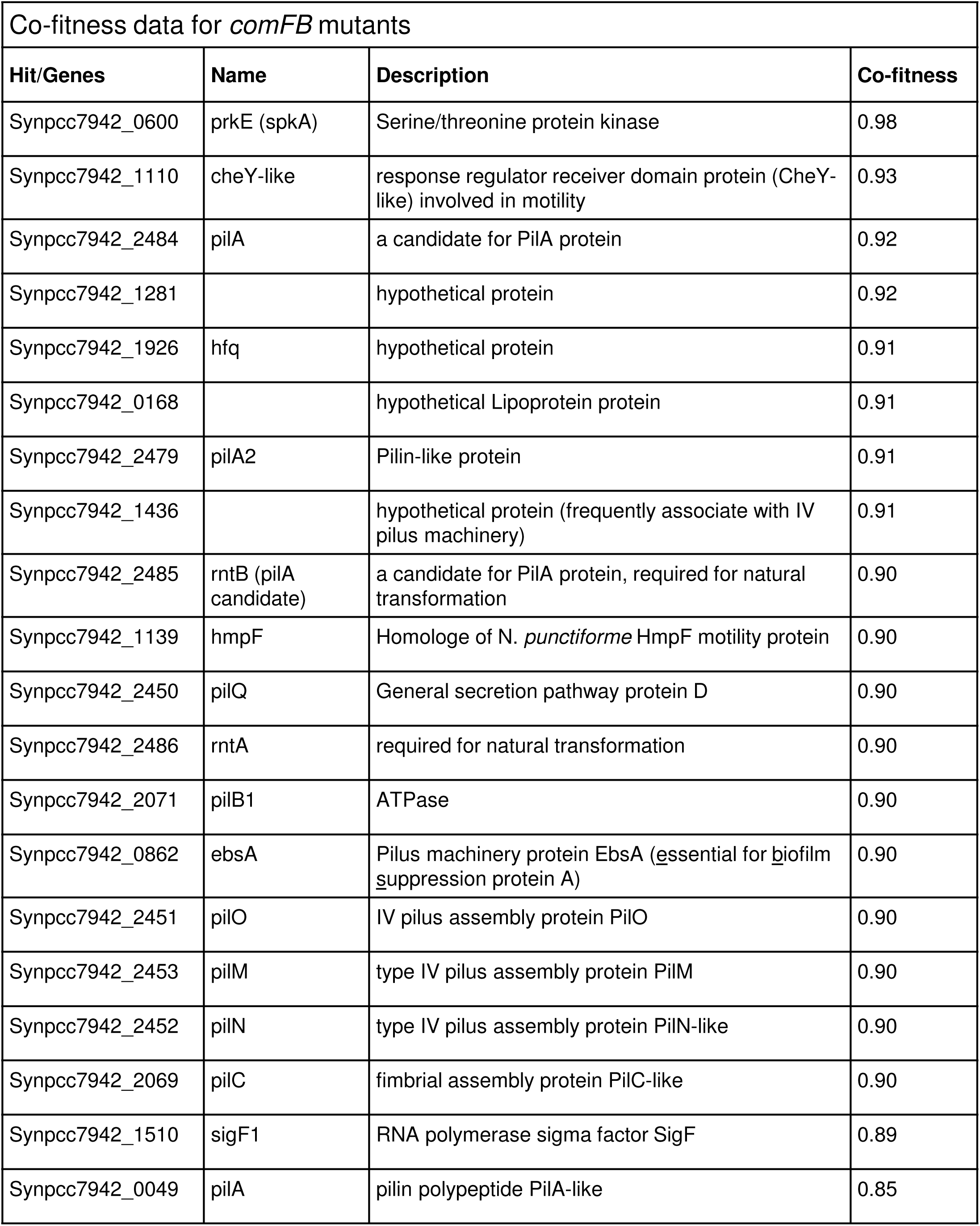
Co-fitness data for *Synechococcus elongatus* Δ*comFB* mutant.

**Fig. S16:**
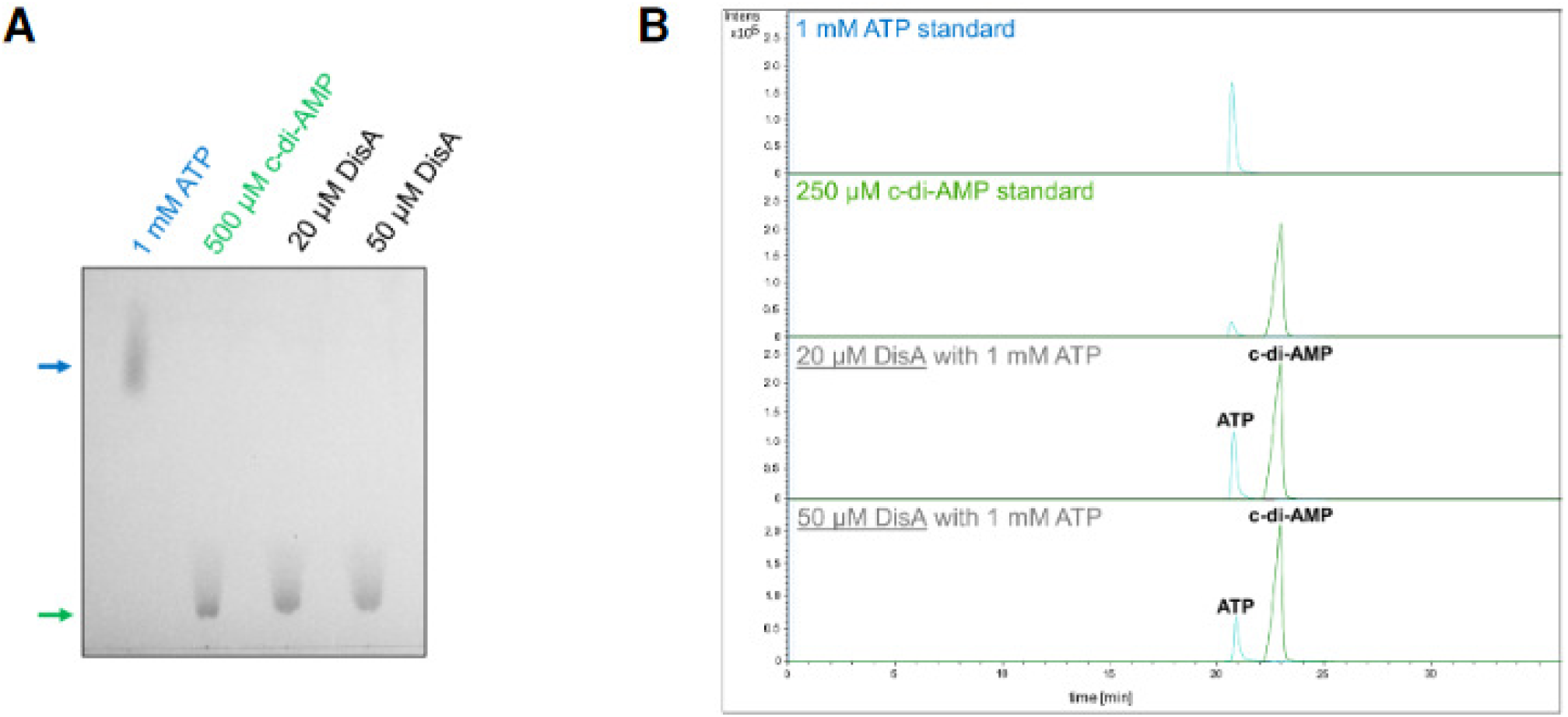
Synthesis of c-di-AMP from purified DisA. Analysis of the enzymatic conversion of 1 mM ATP to c-di-AMP with 20 μM or 50 μM recombinant DisA. (A) Thin layer chromatography (TLC). Arrows denote running distance of c-di-AMP (green) and ATP (blue). (B) LC-MS analysis of purified His_6_-tagged DisA showing conversion of ATP to c-di-AMP.

**Fig. S17:**
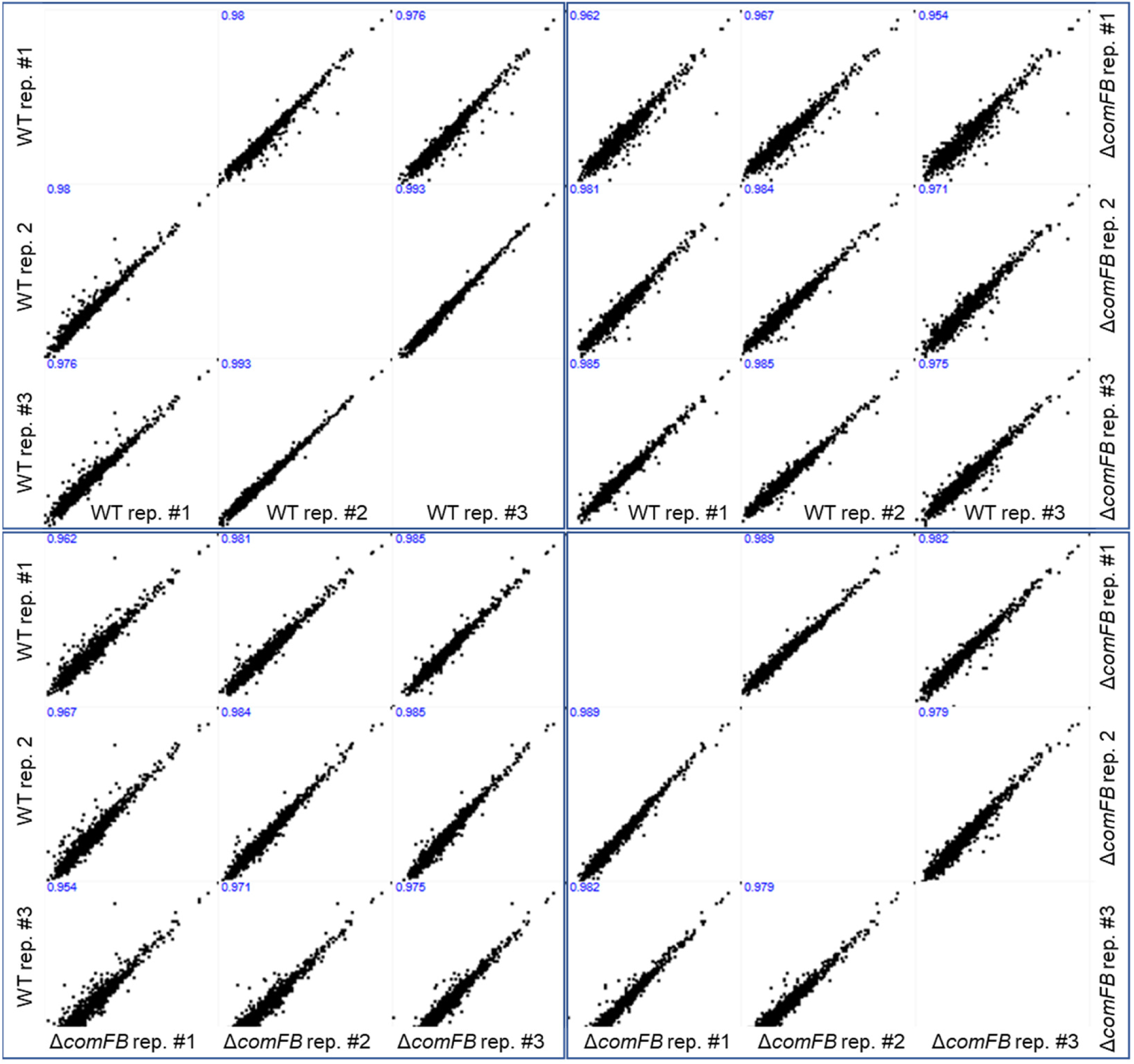
Intensity-based correlation of replicated proteome analyses. Shown is the correlation of protein intensities (in Log_10_ scale) between independent replicates the wildtype (WT) and Δ*comFB* strains. Pearson correlation coefficients are indicated for correlations between each two replicates within the strain and between different strains.

**Fig. S18:**
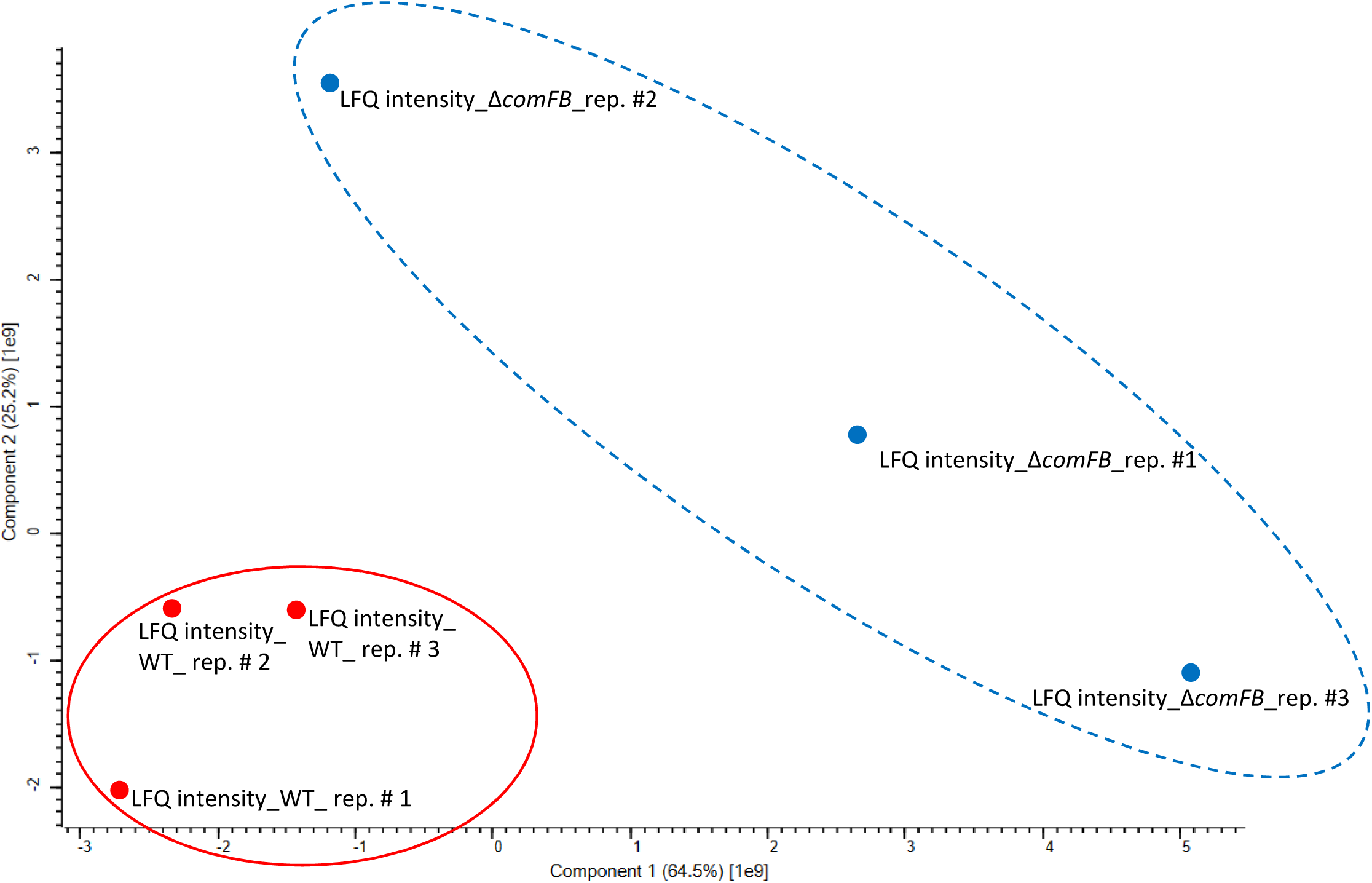
Proteomic landscape of Δ*comFB* mutant in comparison to *Synechocystis* wildtype (WT) cells. Principal component analysis of protein abundance in the wildtype (red circles) and Δ*comFB* (blue circles) is shown. Co-clustering of wildtype replicates indicates similar proteome compositions. Offset clustering of Δ*comFB* (dotted ellipse) indicates broad changes in protein abundance compared to wildtype.

